# Brainstem neurons coordinate the bladder and urethral sphincter for urination

**DOI:** 10.1101/2024.10.01.616107

**Authors:** Xing Li, Xianping Li, Jun Li, Han Qin, Shanshan Liang, Jun Li, Tingliang Jian, Xia Wang, Lingxuan Yin, Chunhui Yuan, Xiang Liao, Hongbo Jia, Xiaowei Chen, Jiwei Yao

## Abstract

Urination, a vital and conserved process of emptying urine from the urinary bladder in mammals, requires precise coordination between the bladder and external urethral sphincter (EUS) that is tightly controlled by a complex neural network. However, the specific subpopulation of neurons that accounts for such coordination remains unidentified, limiting the development of target-specific therapies for certain urination disorders, e.g., detrusor-sphincter dyssynergia. Here, we find that cells expressing estrogen receptor 1 (ESR1^+^) in the pontine micturition center (PMC) initiate voiding when activated and suspend ongoing voiding when suppressed, each at 100% reliability. Transection of the pelvic nerve does not impair PMC^ESR1+^ neurons’ control of the EUS via the pudendal nerve, whereas transection of the pudendal nerve does not impair their control of the bladder via the pelvic nerve. Anatomically, PMC^ESR1+^ neurons consist of three distinct spinal-projection-based subpopulations: one targeting the sacral parasympathetic nucleus (SPN), one innervating the dorsal gray commissure (DGC), and a third that projects to both regions, thereby enforcing the coordination of bladder contraction and sphincter relaxation in a rigid temporal sequence. Thus, we identify a cell type in the brainstem that controls the bladder-urethra coordination for urination.

## Introduction

Urination is a basic life-maintaining function involving the coordinated control of two functional units of the lower urinary tract (Andersson & Anders, 2004; Fowler et al., 2008; Chang et al., 2018), namely the bladder detrusor and external urethral sphincter (EUS). The coordination between the bladder and urethral sphincter for initiating or suspending voiding whenever needed should be executed at 100% reliability in healthy subjects. By contrast, impairment of such coordination (Griffiths et al., 2005; Cho et al., 2015; Taweel & Seyam, 2015), at any rate, leads to various lower urinary tract dysfunctions (Drake et al., 2014; Sakakibara, 2015), significantly degrading the quality of life (Milsom et al., 2014; Aoki et al., 2017). While individual neural control pathways to either the bladder detrusor or the urethral sphincter have been extensively studied (Lee et al., 2021; Xiao et al., 2021), little is known about the neural mechanisms underlying their coordination, which impedes progress in developing target-specific therapies for certain urination disorders, such as detrusor-sphincter dyssynergia (DSD) (Stoffel, 2016; Seseke et al., 2019).

Individual neural pathways are present at different levels of the brain for the control of either the bladder or the EUS (Malykhina, 2017; Jin et al., 2020; Mukhopadhyay & Stowers, 2020). For example, at the cortical level, a subset of motor cortex neurons has been found to drive bladder contraction through the projection to the brainstem (Yao et al., 2018). At the brainstem level, the pontine micturition center (PMC; also referred to as Barrington’s nucleus) has long been considered a command hub region for urination control (Morrison, 2008; Benarroch, 2010). More recent studies have demonstrated that neurons expressing corticotropin-releasing hormone (CRH) in the PMC (PMC^CRH+^) primarily control bladder contraction (Hou et al., 2016; Ito et al., 2020), whereas photostimulation of neurons expressing estrogen receptor 1 (ESR1) in the PMC (PMC^ESR1+^) contributes to sphincter relaxation and increased bladder pressure (Keller et al., 2018). In addition, other subcortical regions, including the periaqueductal gray (PAG) (Rao et al., 2022), lateral hypothalamic area (LHA) (Verstegen et al., 2019; Hyun et al., 2021), and medial preoptic area (MPOA) (Hou et al., 2016), have been suggested to play a potential role in urination control by sending direct projections to the PMC. However, the brain areas that are essential for the coordinated control of both the bladder and the urethral sphincter during reflex voiding remain unclear.

The most likely candidate region for coordinated control is the PMC. Neural tracing experiments demonstrate that the PMC directly sends two bundles of glutamatergic axonal projections, one to the sacral parasympathetic nucleus (SPN), where parasympathetic bladder motoneurons are located, which send axons through the pelvic nerves for bladder control; and the other one to the lumbosacral dorsal gray commissure (DGC) interneurons, which inhibit the sphincter motoneurons in the dorsolateral nucleus (DL) for sphincter control via the pudendal nerves (Jin et al., 2020; Kawatani et al., 2021). Early studies show that microinjection of drugs into the PMC or electrical stimulation in the PMC causes bladder contraction, sphincter relaxation, and urination (Noto et al., 1989; Mallory, 1991; Sugaya & De Groat, 1994). However, the PMC consists of molecularly disparate cell subtypes characterized by the expression of different marker genes, e.g. CRH positive cells (Ito et al., 2020; Van Batavia et al., 2021), ESR1 positive cells (Vanderhorst et al., 2005), vesicular glutamate transporter (Vglut2) and vesicular GABA transporter (VGAT) positive cells (Hou et al., 2016; Verstegen et al., 2017). What remains unknown to date is which exact neuronal subpopulation in the PMC accounts for such coordination of the bladder and urethral sphincter. As previous anatomical results have shown that PMC^CRH+^ neurons mainly project to the SPN in the spinal cord, whereas PMC^ESR1+^ neurons project to both the SPN and DGC in the spinal cord (Keller et al., 2018; Kawatani et al., 2021), we hypothesize that PMC^ESR1+^ cells could be a candidate for the coordination control and began our investigation.

## Results

### Voiding tightly correlates with PMC cell activity

We performed fiber photometry monitoring of neuronal population Ca^2+^ activity (Rao et al., 2022) (fluorescence indicator: GCaMP6f) in the PMC of awake, unrestrained ESR1-Cre mice (Figure supplements 1A and 1B, see Methods for detail). Each detected voiding event tightly correlated with a detected Ca^2+^ transient (Ca^2+^ transient preceding voiding by 0.7\0.3-0.9 s, median\25%-75% percentile, same notation hereinafter; n = 260 voiding events from 9 mice; Figure supplements 1C and 1D), as verified by both an analysis of temporally shuffled data (peak Δf/f value, data: 31.8%\26.7%-33.2%; shuffled: 5.4%\3.7%-6.0%, n = 260 events of 9 mice, *p* = 3.9e-3, Wilcoxon signed-rank test; Figure supplement 1E) and a blank control with an expression of inert fluorescence indicator, EYFP (Enhanced Yellow Fluorescent Protein), in PMC^ESR1+^ cells (detected voiding-associated fluorescence signal events: 100%\100% – 100%, n = 9 mice in test group; 0%\0% – 0%, n = 9 mice in control group, *p* = 4.1e-5, Wilcoxon rank-sum test; Figure supplements 1F and 2).

To further test single-neuron correlates of voiding, we performed an ‘opto-tagging’ experiment (Qin et al., 2022) with a tetrode-fiber bundle implanted in the PMC of ESR1-Cre mice injected with AAV2/8-DIO-ChR2-mCherry (Figure 1A; see Methods for detail). Single PMC^ESR1+^ units (PMC^ESR1+^ cells) were sorted and tagged by detecting reliable spikes to brief optical stimulation (e.g., latency: 4.04\3.02-4.6 ms, success rate: 97.5%\91.5%-100%; Figure supplements 3A-3E). Among 11 PMC^ESR1+^ units that showed urination-related excitation, 8 units exhibited a consistent firing increase in every voiding trial, whereas the remaining 3 increased their discharge in >78 % of trials (Figure 1B and Figure supplement 3F). In contrast, other opto-tagged units did not show increased firing during urination (n = 17 cells; Figure 1B and Figure supplement 3G), indicating functional heterogeneity within the PMC^ESR1+^ population and suggesting that these neurons may participate in other pelvic-related functions (Rouzade-Dominguez et al., 2003; Schellino et al., 2020; Quaghebeur et al., 2021). Importantly, the baseline-corrected firing rate of PMC^ESR1+^ units significantly exceeded that of non-PMC^ESR1+^ units in the temporal association window with voiding (PMC^ESR1+^ units: ‘before’, 0.04\-0.5-0.5 Hz, ‘voiding’, 1.4\-2.5e-11-3.8 Hz, n = 28 cells, *p* = 0.004; non-PMC^ESR1+^ units: ‘before’, 0.8\0.2-1.5 Hz, ‘voiding’, 1\-0.9-1.9 Hz, n = 51 cells, *p* = 0.6; Wilcoxon signed-rank test; Figures 1B and 1C). Overall, these results suggest that PMC^ESR1+^ cells’ firing activities are tightly associated with voiding.

**Figure 1.**
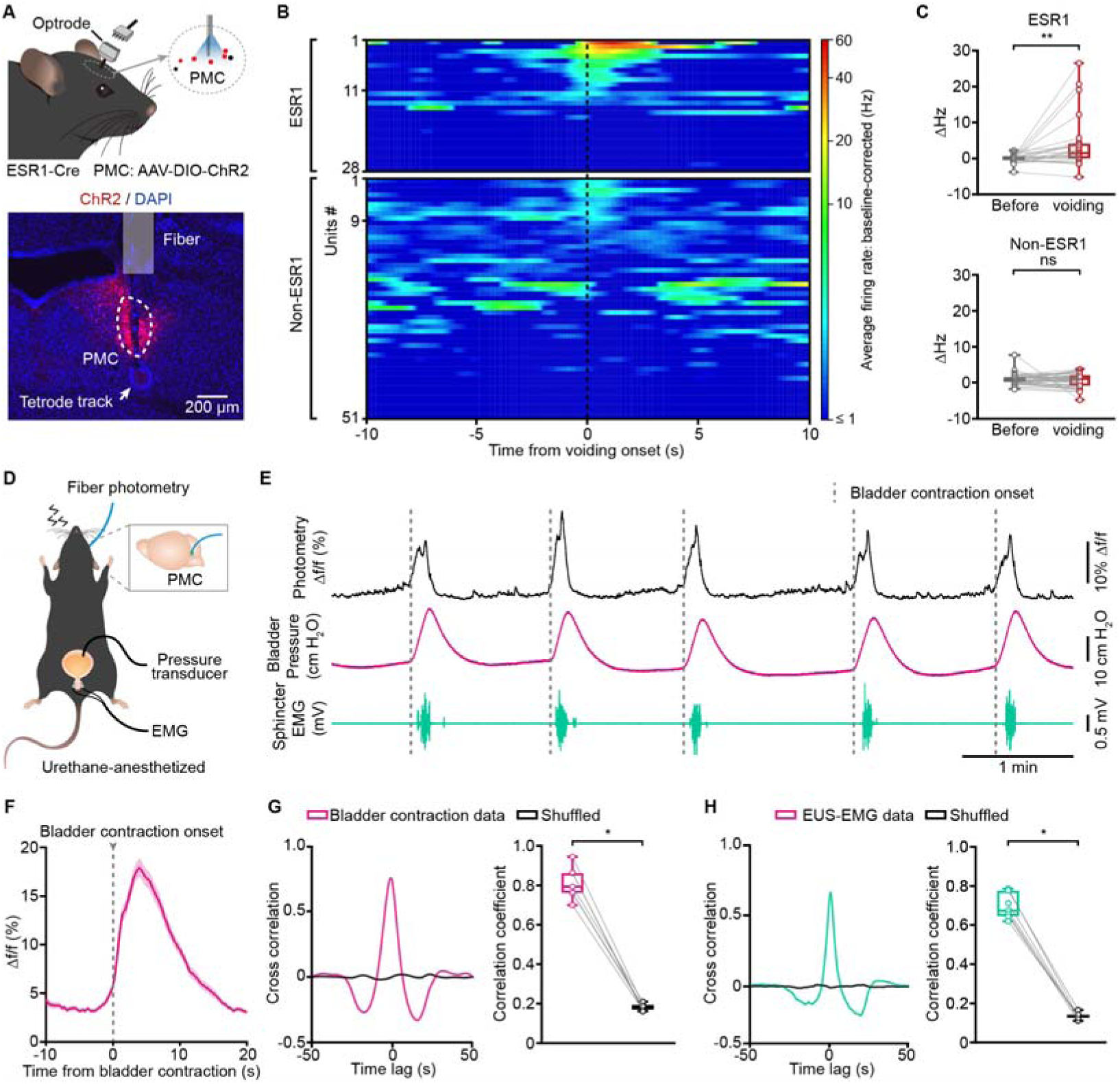
The activity of PMC cells tightly correlates with bladder contraction and sphincter relaxation during successful voiding. (A) Schematic (top) and representative histology (bottom) of optrode recording in the PMC of an ESR1-cre mouse. Scale bar: 200 µm. (B) Cumulative sessions of sorted single-unit activity of PMC^ESR1+^ (upper; n = 28 cells from 4 mice) and non-PMC^ESR1+^ cells (lower; n = 51 cells from 4 mice) aligned to voiding onset (black dashed line), vertically arranged by their instantaneous firing rate at the voiding onset. (C) Boxplots showing the baseline-corrected average firing rates before and during voiding among PMC^ESR1+^ (top, n = 28 cells from 4 mice, \*\**P* = 0.004) and non-PMC^ESR1+^ cells (bottom, n = 51 cells from 4 mice, *P* = 0.6; n.s., not significant; Wilcoxon signed-rank test). (D) Schematic of fiber photometry recording for PMC^ESR1+^ cells during simultaneous cystometry and urethral electromyography in a urethane-anesthetized mouse. (E) Representative traces showing Ca^2+^ transients (black), bladder pressure (magenta), and EUS-EMG (teal) during fiber photometry recordings, with dashed lines indicating bladder contraction onset. (F) Average Ca^2+^ signals during bladder contraction from all trials (n = 101 trials from 7 mice). The thick line and shading represent mean ± s.e.m., respectively. (G) Cross-correlation (left) and correlation coefficients (right) between Ca^2+^ signals and bladder contraction events compared to shuffled data (n = 7 mice, \**P* = 0.02, Wilcoxon signed-rank test). (H) Cross-correlation (left) and correlation coefficients (right) between Ca^2+^ signals and EUS-EMG bursting events compared to shuffled data (n = 7 mice, \**P* = 0.02, Wilcoxon signed-rank test). For all data points in (C, G), and (H), whisker-box plots indicate the median with the 25%-75% percentile as the box, and whiskers represent the minimum and maximum values.

Additionally, we performed a set of combined physiological monitoring experiments, integrating fiber photometry, cystometry, and electromyography (EMG) of external urethral sphincter (EUS) simultaneously in urethane-anesthetized mice (Figure 1D, see methods for detail). For quantitative measurement, we applied cystometry by continuously infusing saline into the bladder through a microcatheter (30-50 μl/min) to induce regular reflexive voiding. This protocol produced cyclic rises in intravesical pressure that were accompanied by stereotyped EUS-EMG bursting (Figure 1E). The EUS bursting activity consists of high-amplitude, high-frequency spike clusters (active periods) interspersed with low tonic activity (silent periods) and generates rhythmic sphincter contractions and relaxations. The silent phases correspond to sphincter relaxation windows that allow urine passage (Kadekawa et al., 2016). Consequently, voiding occurred during the relaxation intervals and was followed by a prompt pressure drop. A triple correlation of events was routinely observed, i.e., PMC^ESR1+^ neuronal activity, bladder pressure elevation, and the bursting pattern of EUS-EMG that cyclically relaxes the sphincter (n = 7 mice, *p* = 0.02, Wilcoxon signed-rank test; Figures 1F-1H). This result was further validated by shuffled data analysis (Figures 1G and 1H), demonstrating the robustness and time-locking precision of the triple correlation events.

However, it is important to note that the association between PMC^ESR1+^ cell activity and voiding in the other way around was not always 100%, i.e., for each detected Ca^2+^ transient, there could be either a voiding contraction (VC) or occasionally a non-voiding contraction (NVC) (Biallosterski et al., 2011) (Figure supplements 4A, 4C and 4D). Nevertheless, not only was the peak of bladder pressure lower (Figure supplement 4E), but also the amplitude of the photometry Ca^2+^ transient, which is known as a reliable report of collective neuronal population activity level, was significantly lower in NVC events than in VC events (NVCs: 8.8%\6.6%-14.7%, n = 62 events from 3 mice; VCs: 13.1%\10.7%-21.8%, n = 79 events from 3 mice, *p* = 8.2e-7, Wilcoxon rank-sum test; Figure supplement 4F), suggesting that not only the timing but also the strength of PMC^ESR1+^ cell activities were tightly correlated with successful voiding. Furthermore, after aligning the fiber-photometry traces to the onset and offset of each EUS bursting episode, a small but consistent hump in the Ca²⁺ signal appeared before bursting onset and the Ca²⁺ signal continued to rise throughout the bursting (Figure supplement 4B, yellow arrow). The Ca²⁺ amplitude at bursting offset was significantly higher than both the NVC peak and the level recorded at bursting onset, (NVCs: 62 events from 3 mice, VCs: 79 events from 3 mice; ***P = 4.4e-4, NVC peak versus VC bursting offset, Wilcoxon rank-sum test; ***P = 1.1e-14, VC bursting onset versus VC bursting offset, Wilcoxon signed-rank test; Figure supplement 4F), indicating that urethral fluid flow/activation supplies excitatory feedback that reinforces PMC activity and bladder contraction, thereby facilitating successful voiding in accordance with Barrington’s classic reflex (Barrington, 1921; Sasaki, 2004).

### PMC cells bidirectionally operate the bladder and sphincter to initiate or suspend voiding

With the tight correlation established, we moved on to test the causal relation between PMC^ESR1+^ cell activity and voiding. We started with a ‘loss-of-function’ test in awake mice, i.e., acute photoinhibition of PMC^ESR1+^ cells in a manual closed-loop to trigger the photoinhibition light as soon as the first patch of urine visualized (Figure 2A and Figure supplement 5A; see Methods for detail). This test was also accompanied by a blank control in which all experimental conditions were the same except that the inhibitory opsin GtACR1 was absent. In photoinhibition events but not in the blank control events, the urine spot area was significantly reduced (‘Pre’: 34.1\27.1-40.4 cm^2^; ‘On’: 9.9\7.9-11.1 cm^2^, n = 12 mice in test group, *p* = 4.9e-4; ‘Pre’: 35.2\27.9-43.9 cm^2^; ‘On’: 38.9\29.98-43.6 cm^2^, n = 8 mice in control group, *p* = 0.3; Wilcoxon signed-rank test; Figures 2B and 2C), and the urination duration was significantly reduced as well (‘Pre’: 5.6\5.1-6.6 s; ‘On’: 1.4\1.2-1.4 s, *p* = 4.9e-4, Wilcoxon signed-rank test; Figures 2D and 2E). Effectively, the ongoing urination was fully suspended after a latency of 0.3 ± 0.1 sec from the onset of photoinhibition (Figure supplement 5B).

**Figure 2.**
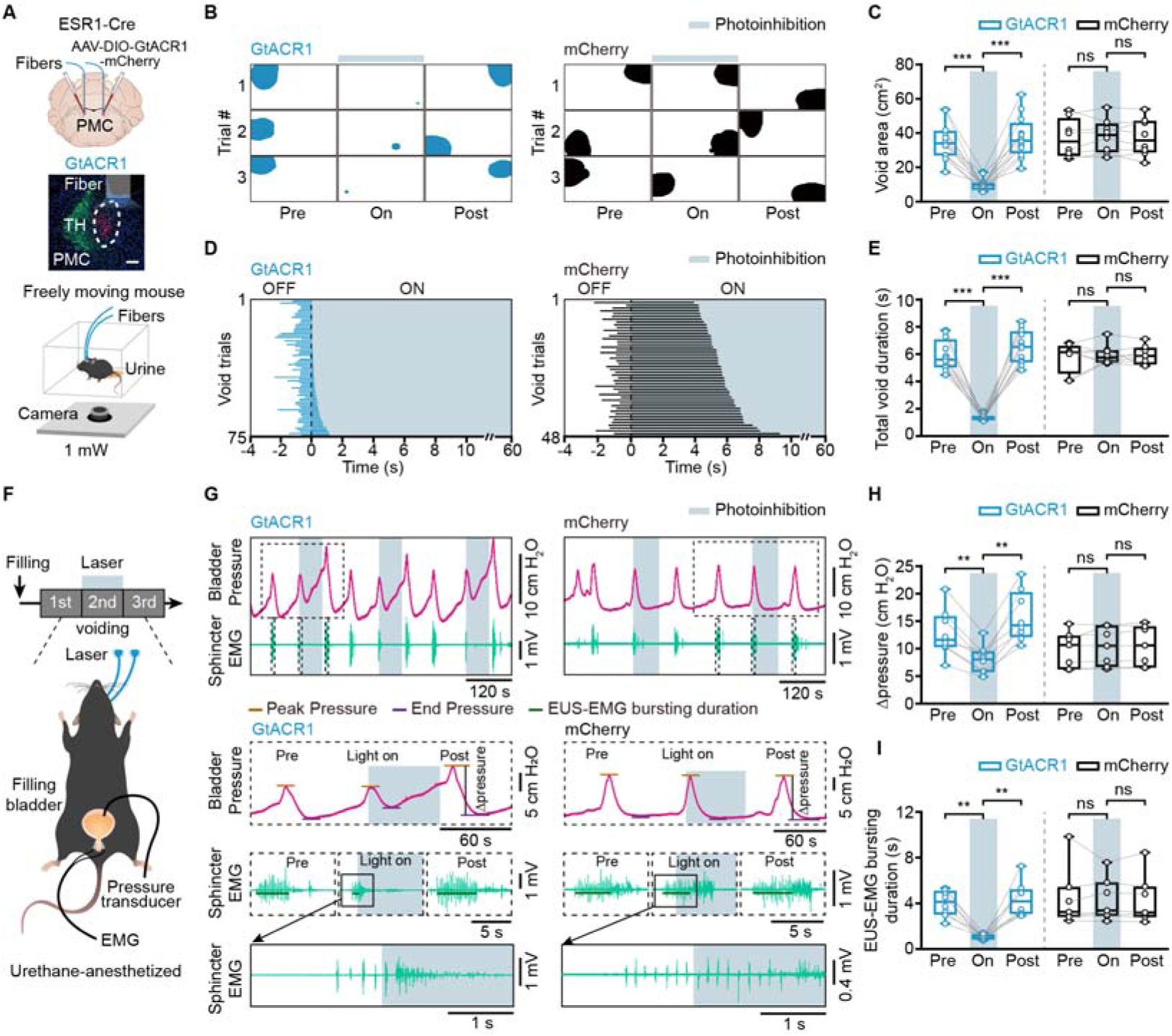
Inactivation of PMC cells suppresses bladder contraction and EUS bursting activity to suspend voiding. (A) Schematic of labeling (top), representative histology (middle), and behavior test (bottom) for PMC^ESR1+^ photoinhibition. Scale bar: 100 µm. (B, C) Representative images (B) and quantification (C) of the void area before (‘Pre’), during (‘On’), and after (‘Post’) photoinhibition in PMC^ESR1-GtACR1^ (n = 12 mice) and PMC^ESR1-mCherry^ (n = 8 mice) groups (from left to right: \*\*\**P* = 4.9e-4, \*\*\**P* = 4.9e-4, *P* = 0.3, *P* = 0.5, respectively; n.s., not significant; Wilcoxon signed-rank test). (D) Cumulative trials of voiding duration in PMC^ESR1-GtACR1^ (blue bar, n = 75 trials from 12 mice) and PMC^ESR1-mCherry^ (black bar, n = 48 trials from 8 mice) during photoinhibition. Voiding trials are ordered by the increasing time of the voiding epoch with the laser on. (E) Voiding duration before, during, and after photoinhibition in PMC^ESR1-GtACR1^and PMC^ESR1-mCherry^ groups (from left to right: \*\*\**P* = 4.9e-4, \*\*\**P* = 4.9e-4, *P* = 0.7, *P* = 0.9, respectively; n.s., not significant; Wilcoxon signed-rank test). (F) Timeline (top) and schematic (bottom) for PMC^ESR1+^ cells photoinhibition during simultaneous cystometry and electromyography recording. (G) Representative traces (top) and expanded portions (bottom) of bladder pressure (magenta) and EUS-EMG (teal) before, during, and after photoinhibition in PMC^ESR1-GtACR1^ (left) and PMC^ESR1-mCherry^ (right) groups. (H, I) Quantification of the Δpressure (H) and EUS-EMG bursting duration (I) during voiding before, during, and after photoinhibition in PMC^ESR1-GtACR1^ (n = 8 mice) and PMC^ESR1-mCherry^ (n = 7 mice) groups (H: from left to right: \*\**P* = 7.8e-3, \*\**P* = 7.8e-3, P = 0.2, *P* = 0.3, respectively; I: from left to right: \*\**P* = 7.8e-3, \*\**P* = 7.8e-3, *P* = 0.8, *P* = 0.4, respectively; n.s., not significant; Wilcoxon signed-rank test). For all data points in (C, E, H), and (I), whisker-box plots indicate the median with the 25%-75% percentile as the box, and whiskers represent the minimum and maximum values.

To understand the physiological process during photoinhibition of PMC^ESR1+^ cells, we performed a simultaneous recording by measuring both the bladder pressure and the electromyograph of the external urethral sphincter (EUS-EMG) under urethane anesthesia (Figure 2F; see Methods for detail). This experiment also involved a manual closed-loop operation to trigger the light when observing the onset of the phasic bursting activity of the EUS-EMG that is known to be directly associated with successful voiding in rodents (Kadekawa et al., 2016; Langdale & Grill, 2016). The maximum relative change in bladder pressure (ΔPressure) upon voiding event was significantly reduced during photoinhibition compared to pre-inhibition baseline events (‘Pre’: 11.6\10.4-15.4 cm H_2_O, ‘On’: 8.1\5.9-9.4 cm H_2_O, n = 8 mice, *p* = 7.8e-3, Wilcoxon signed-rank test), but no such reduction was observed in the blank control group (‘Pre’: 10.7\6.9-12.03 cm H_2_O, ‘On’: 10.5\7.2-13.4 cm H_2_O, n = 7 mice, *p* = 0.2, Wilcoxon signed-rank test; Figures 2G and 2H). In the meanwhile, photoinhibition of the PMC^ESR1+^ cells also halted the voiding-associated, phasic bursting activity of EMG, transform to a tonic activity, effectively reducing the sphincter bursting duration (‘Pre’: 4.1\3.2-4.9 s, ‘On’: 0.98\0.9-1.2 s, *p* = 7.8e-3, Wilcoxon signed-rank test; Figures 2G and 2I) at a latency of 0.1 ± 0.02 sec (Figure supplements 5C and 5D), but no such effect was observed in the blank control group (‘Pre’: 3.3\2.9-4.8 s, ‘On’: 3.4\2.96-5.4 s, *p* = 0.8, Wilcoxon signed-rank test). Furthermore, the immediate suspension of an ongoing voiding event by photoinhibition of PMC^ESR1+^ cells resulted in an expected side effect that the threshold of bladder pressure to initiate the next voiding event (post-photoinhibition) became higher (Figure supplement 5E). A control test for the above set of ‘loss-of-function’ experiments was to test whether shorter durations of photoinhibition also had the same urination suspension effect. To address this, we performed additional sets of control experiments in which a shorter duration of photoinhibition (5 s light-on, instead of 60 s) resulted in the same, 100% suspension effect (Figure supplement 6). These data together reveal that the acute photoinhibition of PMC^ESR1+^ cells halted both bladder contraction and the EUS bursting activity that produces sphincter relaxation, thereby leading to a full suspension of the ongoing voiding process.

Then, we performed a ‘gain-of-function’ test in awake mice, i.e., acute photoactivation of PMC^ESR1+^ cells that expressed the excitatory opsin channelrhodopsin-2 (ChR2), in a semi-closed loop to trigger the light on when the bladder was filled to a level that was lower than the threshold required for spontaneous voiding reflex (light on at 3.1 ± 0.03 min after the previous voiding; inter-voiding interval under control condition: 6.6 ± 0.5 min; Figure supplement 7A; see Methods for detail). This test was also accompanied by a blank control in the absence of ChR2. Light stimulation initiated voiding event at 100% reliability (100%\100%-100%, n = 172 trials of 8 mice; Figure supplements 7B and 7C) in the test group (response latency, 0.7\0.6-0.8 s; Figure supplement 7D), but almost 0% (0%\0%-1.25%, n = 166 trials of 8 mice) in the blank control group (Figure supplements 7B and 7C).

Accordingly, we also performed simultaneous cystometry with the EUS-EMG experiment under urethane anesthesia (Figure supplement 7E; see Methods for detail). Light stimulation induced a prominent upstroke of bladder pressure in the test group and did not affect bladder pressure at all in the control group (ΔP: 5.6\4.5-8.2 cm H_2_O, n = 9 mice in the test group, −0.1\-0.2-0.1 cm H_2_O, n = 6 mice in the control group, *p* = 4e-4, Wilcoxon rank-sum test; Figure supplements 7F-7H). Meanwhile, light stimulation triggered the phasic bursting activity of EUS-EMG, at a success rate of 100% in the test group and nearly 0% in the control group (Figure supplements 7F and 7H). An additional set of control experiments by using regular inter-stimulation intervals (5 s light-on per every 30 s) instead of the threshold-adaptive interval, yielded the same 100% success rate of both bladder pressure and EUS-EMG responses (Figure supplement 8). Thus, photoactivation of PMC^ESR1+^ cells initiate voiding by eliciting bladder contraction and triggering EUS bursting, which generates rhythmic sphincter contractions and relaxations. These sets of ‘loss-of-function’ and ‘gain-of-function’ experiments together demonstrate that PMC^ESR1+^ cells perform as a 100% reliable ‘master switch’ for either initiate or suspend voiding, provided that all downstream targets are directly controlled and operational, which will be tested next.

### Transection of either the pudendal or pelvic nerve does not impair PMC neuronal control of the other

To further confirm whether the effect of PMC^ESR1+^ cells on the bladder can occur independently of sphincter relaxation, we designed a new set of simultaneous cystometry and EUS-EMG experiments with PMC^ESR1-ChR2^ mice subjected to first a pudendal nerve transection (PDNx, disrupting the pathway from the PMC to the EUS) and then additionally a pelvic nerve transection (PLNx, disrupting the pathway from the PMC to the bladder) in the same mice under urethane anesthesia (Figure 3A; see Methods for detail). Under PDNx condition, PMC^ESR1+^ cell photoactivation could not elicit EUS-EMG response at all, but robustly elicited bladder pressure upstroke which was then fully abolished after PLNx (Figures 3B-3E). Notably, the photoactivation-induced bladder pressure upstroke under the PDNx condition was nearly the same as that in the pre-transection control condition (Figure 3D), suggesting that the PMC^ESR1+^ cell control of the bladder was fully operational even when the pudendal nerve to sphincter was severed.

**Figure 3.**
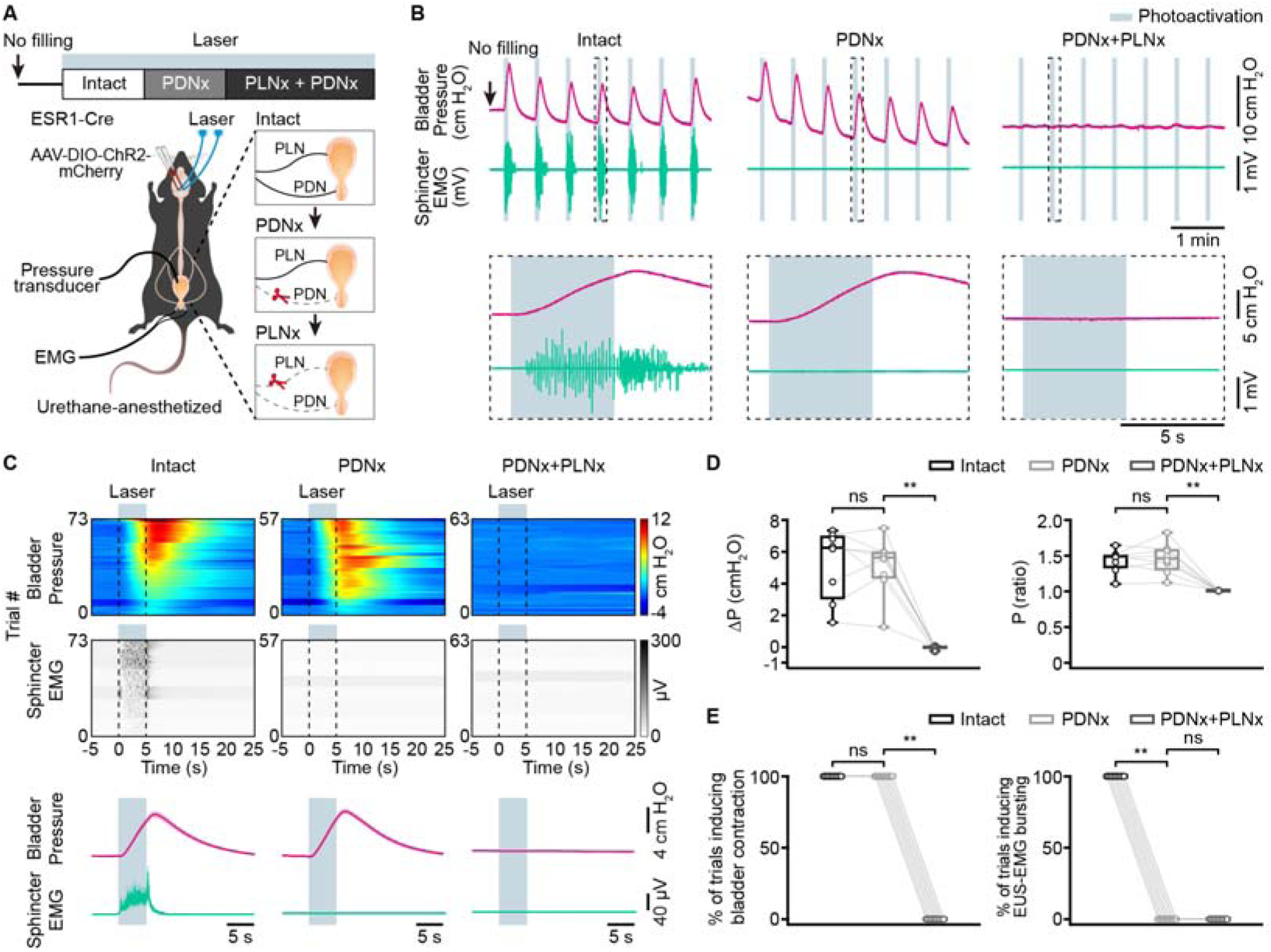
Transection of the pudendal nerves does not impair bladder contraction induced by PMC^ESR1+^ cell photoactivation. (A) Timeline (top) and schematic (bottom) for PMC^ESR1-ChR2^ photoactivation during simultaneous cystometry and urethral electromyography recordings, with PDNx performed first. PDNx, pudendal nerve transection; PLNx, pelvic nerve transection. (B) Representative traces (top) and expanded portions (bottom, from the dashed box in the top panel) showing bladder pressure (magenta) and EUS-EMG (teal) during PMC^ESR1-ChR2^ photoactivation in various groups, with PDNx performed first. (C) Heatmap (top) and average traces (bottom; thick lines and shading represent mean ± s.e.m.) of sorted bladder pressure and EUS-EMG around the photoactivation timepoint for all unfilled bladder trials with the PDNx-first experiment (n = 8 mice per group). (D, E) Quantification of bladder pressure change (ΔP, D, left), bladder pressure ratio (D, right), the percentage of photoactivation-associated bladder contraction (E, left), and the percentage of photoactivation-associated EUS-EMG bursting (E, right) upon photoactivation for the PDNx-first experiment from (C) (n = 8 mice per group, from left to right, D: *P* = 0.8, \*\**P* = 7.8e-3, *P* =0.7, \*\**P* = 7.8e-3, respectively; E: *P* = 1, \*\**P* = 7.8e-3, \*\**P* = 7.8e-3, P = 1, respectively; n.s., not significant; Wilcoxon signed-rank test). For all data points in (D, E), whisker-box plots indicate the median with the 25%-75% percentile as the box, and whiskers represent the minimum and maximum values.

Given that reflex signals from bladder afferents can indirectly influence the urethral sphincter activity (Chang et al., 2007), we next investigated whether activation of PMC^ESR1+^ cells directly induces the bursting pattern of EUS in the absence of bladder contraction. To this end, the sequential nerve transection test was performed again with another group of PMC^ESR1-ChR2^ mice in a different order, i.e., first PLNx to disrupt the bladder nerve and then PDNx to additionally disrupt the urethral sphincter nerve (Figure 4A; see Methods for detail). Interestingly, under the PLNx condition with a filled bladder, PMC^ESR1+^ cell photoactivation did not induce a bladder pressure upstroke (Δpressure: 3.2\0.96-4.9 cm H_2_O in pre-transection control condition, −1.2\-1.4 - −0.8 cm H_2_O in PLNx condition, n = 8 mice, *p* = 7.8e-3, Wilcoxon signed-rank test), but the EUS-EMG activity remained responsive, leading to urine leakage which consistently reduced bladder pressure (Figures 4B-4E). Both the inversed bladder response and the EUS-EMG response were then fully abolished after the second transection, PDNx (Figures 4B-4E and Figure supplement 9). Consistently, under the PLNx condition with an unfilled bladder, the EUS-EMG response was present, and the bladder pressure did not change upon PMC^ESR1+^ cell photoactivation (Figure supplement 9). Collectively, these data together indicate that PMC^ESR1+^ cell control of the urethral sphincter was operational even when the pelvic nerve to the bladder was severed.

**Figure 4.**
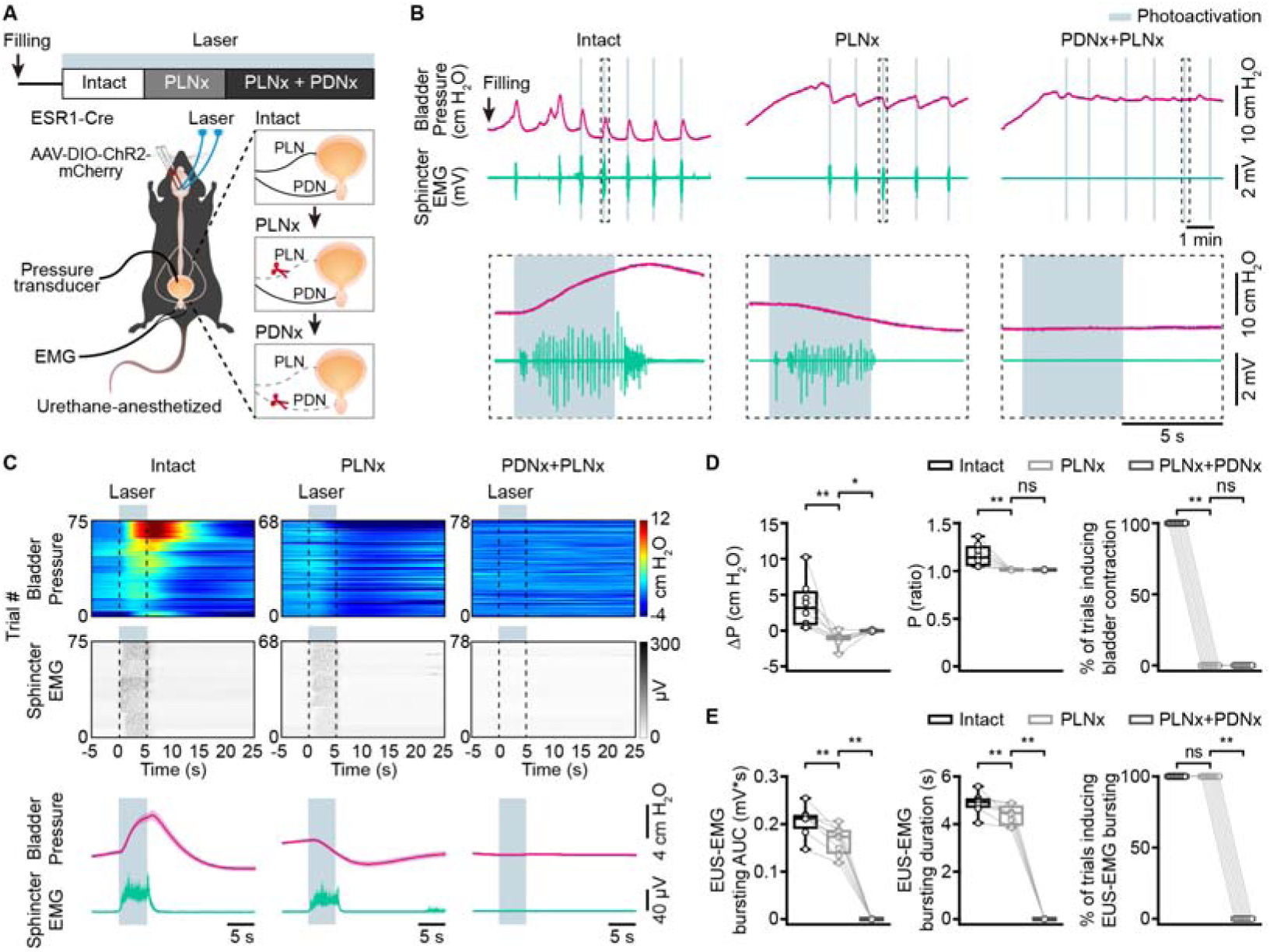
Transection of the pelvic nerves does not impair EUS bursting activity induced by PMC^ESR1+^ cell photoactivation. (A) Timeline (top) and schematic (bottom) for PMC^ESR1-ChR2^ photoactivation during simultaneous cystometry and urethral electromyography recordings, with PLNx performed first. PLNx, pelvic nerve transection; PDNx, pudendal nerve transection. (B) Representative traces (top) and expanded portions (bottom, from the dashed box in the top panel) showing bladder pressure (magenta) and EUS-EMG (teal) during PMC^ESR1-ChR2^ photoactivation in various groups, with PLNx performed first. (C) Heatmap (top) and average traces (bottom; thick lines and shading represent mean ± s.e.m.) of sorted bladder pressure and EUS-EMG around photoactivation timepoint for all filled bladder trials with the PLNx-first experiment (n = 8 mice per group). (D) Quantification of bladder pressure parameters for the PLNx-first experiment from C: bladder pressure change (ΔP, left), bladder pressure ratio (middle), and the percentage of photoactivation-associated bladder contraction (right; from left to right: \*\**P* = 7.8e-3, \**P* = 0.02, \*\**P* = 7.8e-3, *P* = 0.6, \*\**P* = 7.8e-3, *P* = 1, respectively; n.s., not significant; Wilcoxon signed-rank test). (E) Quantification of EUS-EMG parameters for the PLNx-first experiment from C: EUS-EMG bursting AUC (left), EUS-EMG bursting duration (middle), and the percentage of photoactivation-associated EUS-EMG bursting (right; from left to right: \*\**P* = 7.8e-3, \*\**P* = 7.8e-3, \*\**P* = 7.8e-3, \*\**P* = 7.8e-3, *P* = 1, \*\**P* = 7.8e-3, respectively; n.s., not significant; Wilcoxon signed-rank test). For all data points in (D, E), whisker-box plots indicate the median with the 25%-75% percentile as the box, and whiskers represent the minimum and maximum values.

Lastly, despite that the PMC^ESR1+^ cell photoactivation consistently elicited EUS-EMG bursting in 100% of cases, a refined analysis revealed that the key parameters of the sphincter were slightly degraded under PLNx condition (EMG bursting AUC area: 0.21\0.20-0.22 mV*s in pre-transection, 0.17\0.15-0.19 mV*s in PLNx; EMG bursting duration: 4.9\4.8-5.03 s in pre-transection, 4.5\4.1-4.6 s in PLNx, n = 8 mice, *p* = 7.8e-3; Wilcoxon signed-rank test; Figure 4E), which implies a potential reduction in effective voiding volume (Langdale & Grill, 2016). To further explore this, we conducted experiments on awake, unrestrained mice under the PLNx condition (Figure 5A, see Methods for details). Compared to the pre-transection control condition (‘baseline’), PMC^ESR1+^ cell photoactivation still triggered voiding in 100% of trials (Figures 5B and 5C). However, in PLNx mice, the voiding events were characterized by significantly smaller urine spots and a longer latency to voiding onset than the control events before transection, a difference not observed in the sham surgery group (Figures 5D and 5E). These findings could be interpreted as that while the PMC^ESR1+^-EUS pathway is sufficient to drive voiding, as previously reported (Keller et al., 2018), however, without an intact bladder reflex pathway involved, an efficient urine flow cannot be performed.

**Figure 5.**
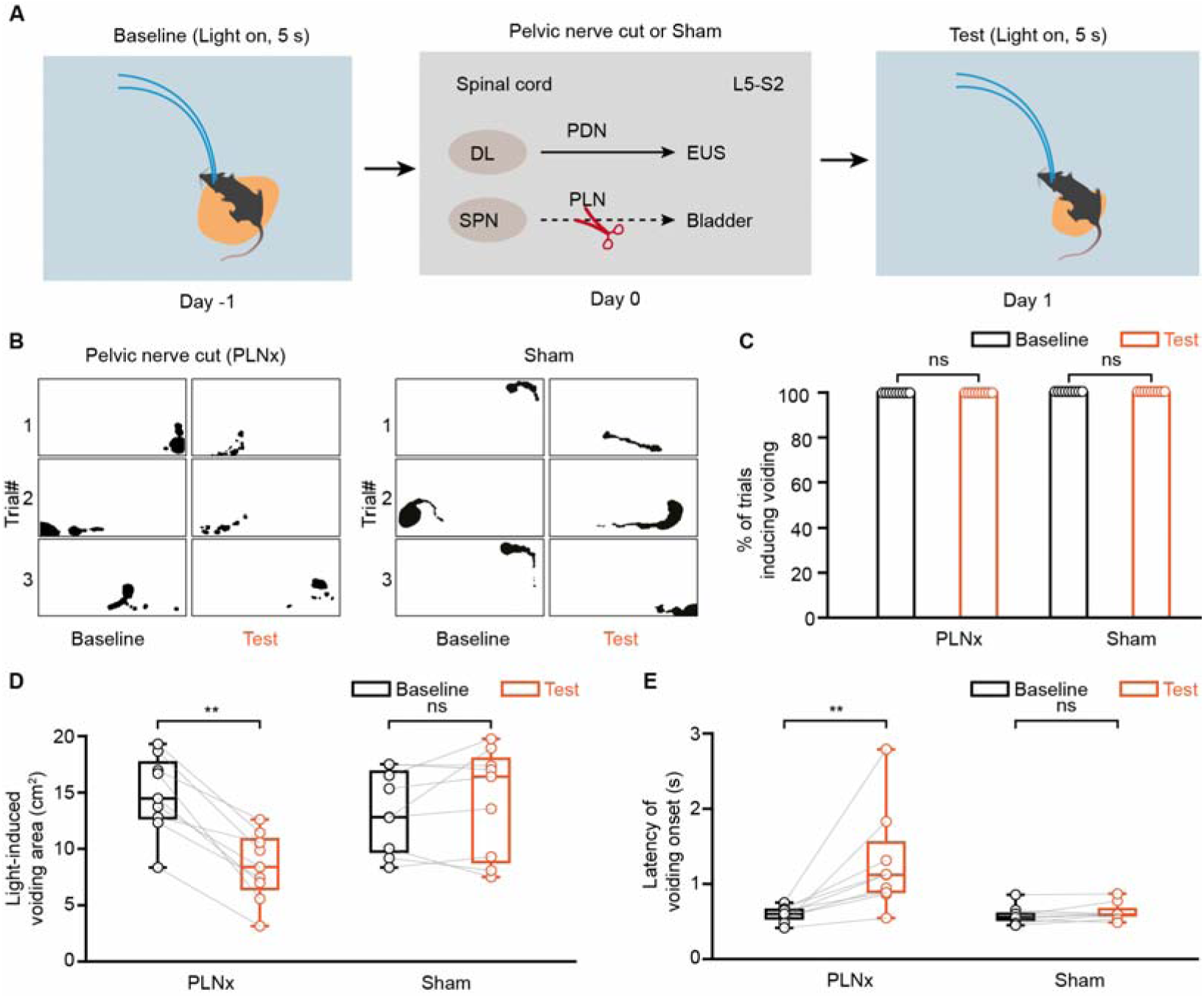
Transection of the pelvic nerves decreases urinary volume induced by PMC cell photoactivation. (A) Schematic (top) and timeline (bottom) for PMC^ESR1-ChR2^ photoactivation in a freely moving mouse with pelvic nerve transection (PLNx). (B) Representative images of light-induced urination marking (black shading) in ESR1-Cre mice before (‘Baseline’) and after (‘Test’) pelvic nerve transection (left) or sham surgery (right). (C-E) Quantification of the effect of pelvic nerve transection on voiding in the PLNx (n = 9 mice) and sham groups (n = 9 mice): the percentage of light-induced voiding (C, P = 1; n.s., not significant; Wilcoxon signed-rank test), light-induced voiding area (D, \*\**P* = 3.9e-3, *P* = 0.5, respectively), and latency of voiding onset after light stimulation (E, \*\**P* = 3.9e-3, *P* = 0.2, respectively; n.s., not significant; Wilcoxon signed-rank test). For all data points in (D, E), whisker-box plots indicate the median with the 25%-75% percentile as the box, and whiskers represent the minimum and maximum values.

Similarly, in awake, unrestrained PDNx mice (Figure 6A), photoactivation of PMC^ESR1+^ cells reliably triggered voiding in all trials (Figures 6B and 6C), but the urine spot area was reduced and the latency to voiding onset did not differ significantly from that of the sham surgery group (Figures 6D and 6E), indicating that effective urine flow still depends on bilateral integrity of the pudendal nerve to entrain rhythmic EUS bursting. Piecing these data of the combined transection-optogenetics experiments together, we reveal a more complete picture that PMC^ESR1+^ cells can operate the bladder (through the pelvic nerve) and the sphincter (through the pudendal nerve) independently of each other. Such refined knowledge could not have been obtained from simpler tests using optogenetics alone as shown above (Figure 2 and Figure supplement 7), or from other experiments in the literature (Hou et al., 2016; Keller et al., 2018; Ito et al., 2020).

**Figure 6.**
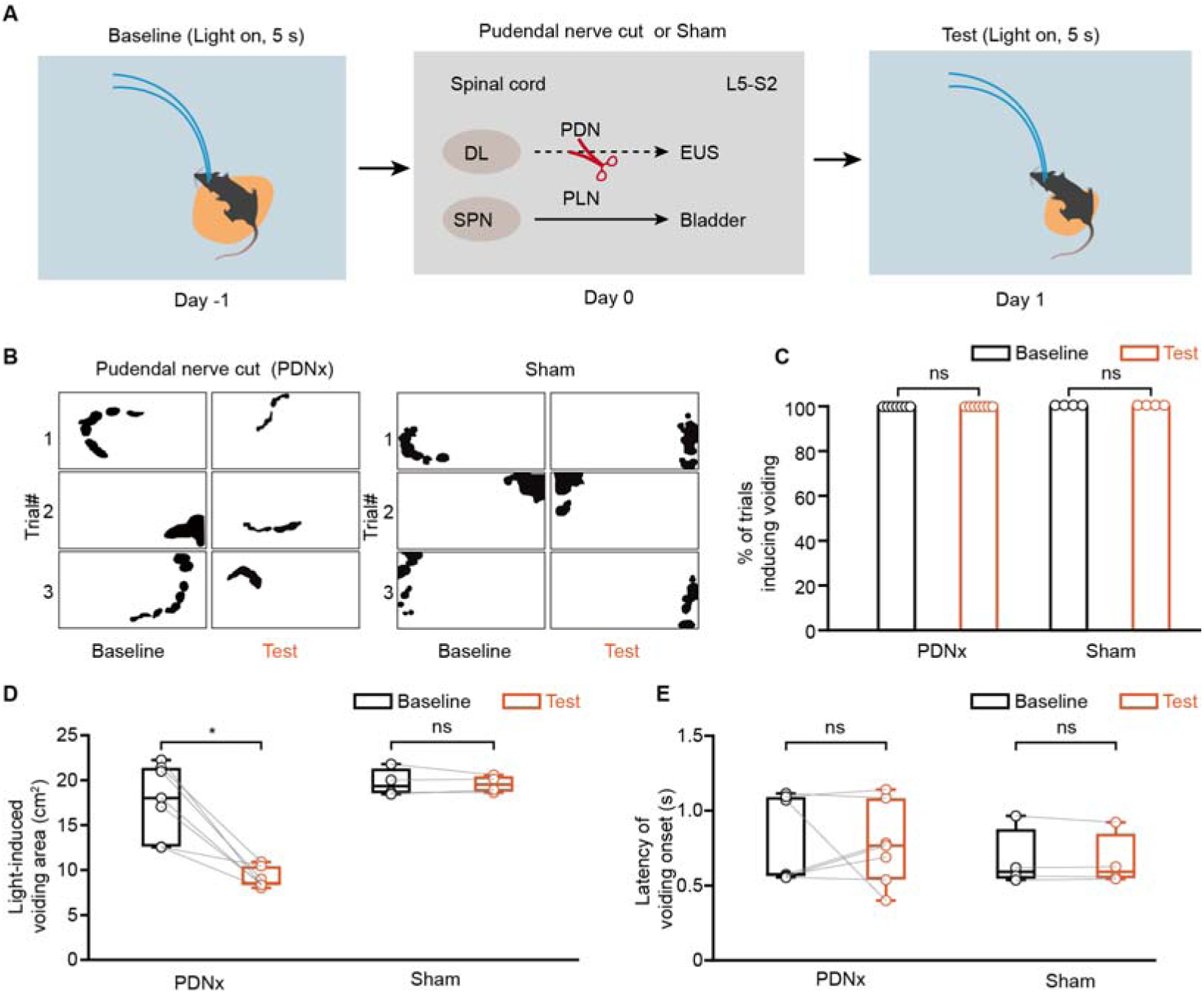
Transection of the pudendal nerves decreases urinary volume induced by PMC^ESR1+^ cell photoactivation. (A) Schematic (top) and timeline (bottom) for PMC^ESR1-ChR2^ photoactivation in a freely moving mouse with pudendal nerve transection (PDNx). (B) Representative images of light-induced urination marking (black shading) in ESR1-Cre mice before (‘Baseline’) and after (‘Test’) pudendal nerve transection (left) or sham surgery (right). (C-E) Quantification of the effect of pudendal nerve transection on voiding in the PDNx (n = 7 mice) and sham groups (n = 4 mice): the percentage of light-induced voiding (C, *P* = 1; n.s., not significant; Wilcoxon signed-rank test), light-induced voiding area (D, \**P* = 1.6e-2, *P* = 1, respectively), and latency of voiding onset after light stimulation (E, *P* = 0.6, P = 0.9, respectively; n.s., not significant; Wilcoxon signed-rank test).For all data points in (D, E), whisker-box plots indicate the median with the 25%-75% percentile as the box, and whiskers represent the minimum and maximum values.

PMC^ESR1+^ cells coordinate bladder contraction and sphincter relaxation to initiate urination. The above data showing that PMC^ESR1+^ cells can operate through both the bladder and urethral sphincter independently of each other implies that there should be two distinct anatomical projections from PMC^ESR1+^ cells downstream to innervate these targets. Indeed, our anterograde labeling experiment in ESR1-Cre mice (Figure supplement 10; see Methods for detail) showed that the mGFP-labelled axonal terminals of the PMC^ESR1+^ cell population were found in both the DGC and the SPN region of the lumbosacral spinal cord. Consistent with these projection targets, DGC interneurons inhibit sphincter motoneurons in Onuf’s nucleus, whose axons project via the pudendal nerve to innervate the urethral sphincter, whereas parasympathetic motoneurons in the SPN send axons via the pelvic nerve to innervate the bladder (Yao et al., 2018; Karnup & De Groat, 2020; Karnup, 2021; Yan et al., 2025).

To further test this hypothesis, we performed retrograde labeling experiments in ESR1-Cre mice and, as a control, in CRH-Cre mice (Figures 7A and 7E; see Methods for detail). Analysis of CRH-Cre mice showed that 80.9% (80.1%\77.3%-80.6%) of PMC^CRH+^ cells were SPN-projecting only (expressing mCherry), and 2.5% (2.1%\1.4%-3.1%) were DGC-projecting only (expressing EGFP), whereas the remaining 16.6% (17.3%\16.8%-18.0%) were dual-projecting (altogether n = 2312 cells pooled from 132 slices obtained from 5 mice; Figures 7B-7D). This result is consistent with the literature (Valentino et al., 2010; Hou et al., 2016; Keller et al., 2018; Ito et al., 2020) that PMC^CRH+^ cells primarily project to the SPN in the spinal cord and modulate bladder contraction. By contrast, analysis of the ESR1-Cre mice showed that 19.0% (20.8%\15.1%-24.1%) of PMC^ESR1+^ cells were SPN-projecting only, 52.2% (50.9%\44.6%-54.4%) were DGC-projecting only, and 28.8% (30.2%\25.9%-31.0%) cells were dual-projecting (n = 2468 cells pooled from 138 slices obtained from 7 mice; Figures 7F-7H). These data suggest that PMC^ESR1+^ cells innervate either SPN or DGC in a more balanced manner and include a significant fraction of dual-innervating cells, in contrast to PMC^CRH+^ cells that primarily innervate the SPN. Consistently, dual-projecting PMC neurons that project SPN and DGC exhibited robust activation throughout the entire voiding phase that was tightly correlated with intravesical pressure rise and EUS bursting (Figure supplements 11A-11H), and optogenetic activation of this population initiates urination with bladder contraction and rhythmic EUS bursting (Figure supplements 11I-11N).

**Figure 7.**
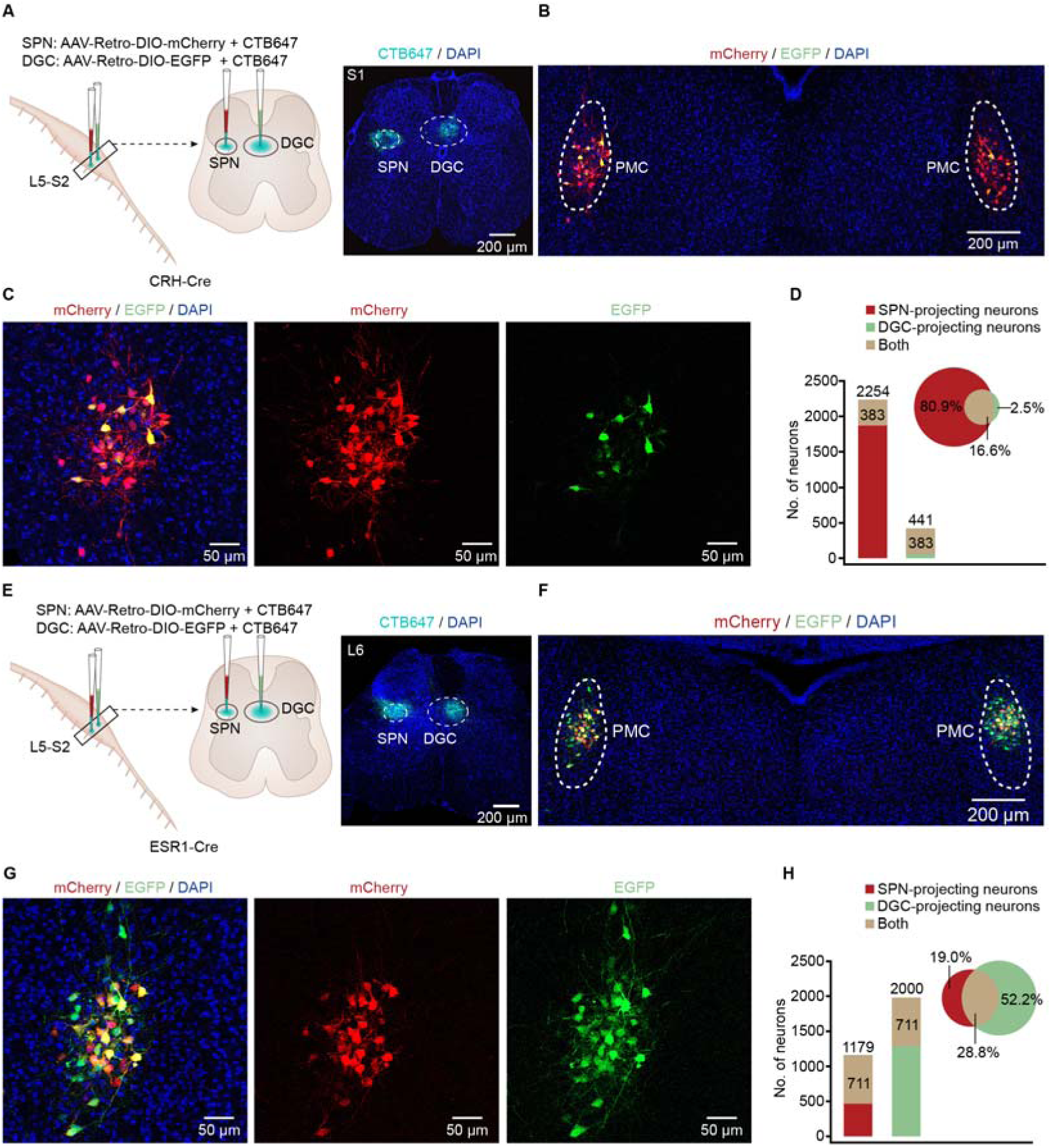
Differences in the anatomical projections to the lumbosacral spinal cord between PMC^CRH+^ and PMC^ESR1+^ cells. (A) Schematic of labeling (left) and representation histology showing the CTB-647 fluorescence in the lumbosacral spinal cord of CRH-Cre mice. Scale bar: 200 µm. (B, C) Representative image (B) and enlarged images (C, from the left part of B) showing EGFP and mCherry expression in the PMC of a CRH-Cre mouse. Scale bars: 200 µm for (B) and 50 µm for (C). (D) Quantification of the fractions of CRH^+^ cells specifically projecting to the SPN and DGC of the spinal cord, respectively (n = 2312 cells from 5 mice). (E) Schematics of labeling (left) and representative histology showing the CTB-647 fluorescence in the lumbosacral spinal cord of ESR1-Cre mice. Scale bar: 200 µm. (F, G) Representative image (F) and enlarged images (G, from the left part of F) showing EGFP and mCherry expression in the PMC of an ESR1-Cre mouse. Scale bars: 200 µm for (F) and 50 µm for (G). (H) Quantification of the fractions of ESR1^+^ cells specifically projecting to the SPN and DGC of the spinal cord, respectively (n = 2468 cells from 7 mice). SPN, sacral parasympathetic nucleus; DGC, dorsal gray commissure.

The functional and anatomical data above together suggest that PMC^ESR1+^ cells can function as a 100% reliable ‘master switch’ either to initiate or to suspend voiding through independently operating the bladder (via SPN to the pelvic nerve) and the sphincter (via DGC to the pudendal nerve). We now come to the final question as to whether PMC^ESR1+^ cells can implement the coordination between the bladder and urethral sphincter, i.e., operate them in a rigid temporal order. We took advantage of the simultaneous recording of cystometry and EUS-EMG condition (e.g., Figure 1) in which the beginning/ending timepoint of bladder pressure upstroke (a significant rapid, transient increase in bladder pressure preceding voiding, denoting threshold pressure of bladder contraction) (Rana et al., 2024) and the stereotypic voiding-associated firing pattern of EUS-EMG could be precisely determined. In the photometry recording experiments when voiding events spontaneously occurred, the bladder pressure upstroke timepoint always preceded the beginning timepoint of the EUS-EMG firing pattern (bladder pressure upstroke onset: 0.6\0.3-1.2 s, EMG bursting onset: 2.3\1.5-3.2 s, relative to the onset of Ca^2+^ signals; n = 46 events pooled from 8 mice; Figure 8A). Accordingly, in the photoactivation experiments, the bladder pressure upstroke timepoint also preceded the EMG bursting pattern begin timepoint, albeit both were slightly earlier than those in ‘passive’ spontaneous photometry recordings (bladder pressure upstroke onset: 0.4\0.4-0.5 s, EMG bursting onset: 0.8\0.7-1.0 s; n = 50 events pooled from 10 mice; Figure 8B). In addition, activation of dual-projecting PMC neurons (to SPN and DGC) consistently led to a sequence in which bladder pressure increased first, followed by rhythmic EUS bursting, during both spontaneous and optogenetic activation induced voiding (Figure supplements 12A and 12B). Despite that the photoactivation of the PMC^ESR1+^ cells could have been artificially strong and did not necessarily mimic the naturalistic firing pattern of PMC^ESR1+^ cells (see Figures 1A-1C), the same temporal order as bladder contraction preceded sphincter relaxation suggests that the downstream circuity was instructed to execute in the same temporal order.

**Figure 8.**
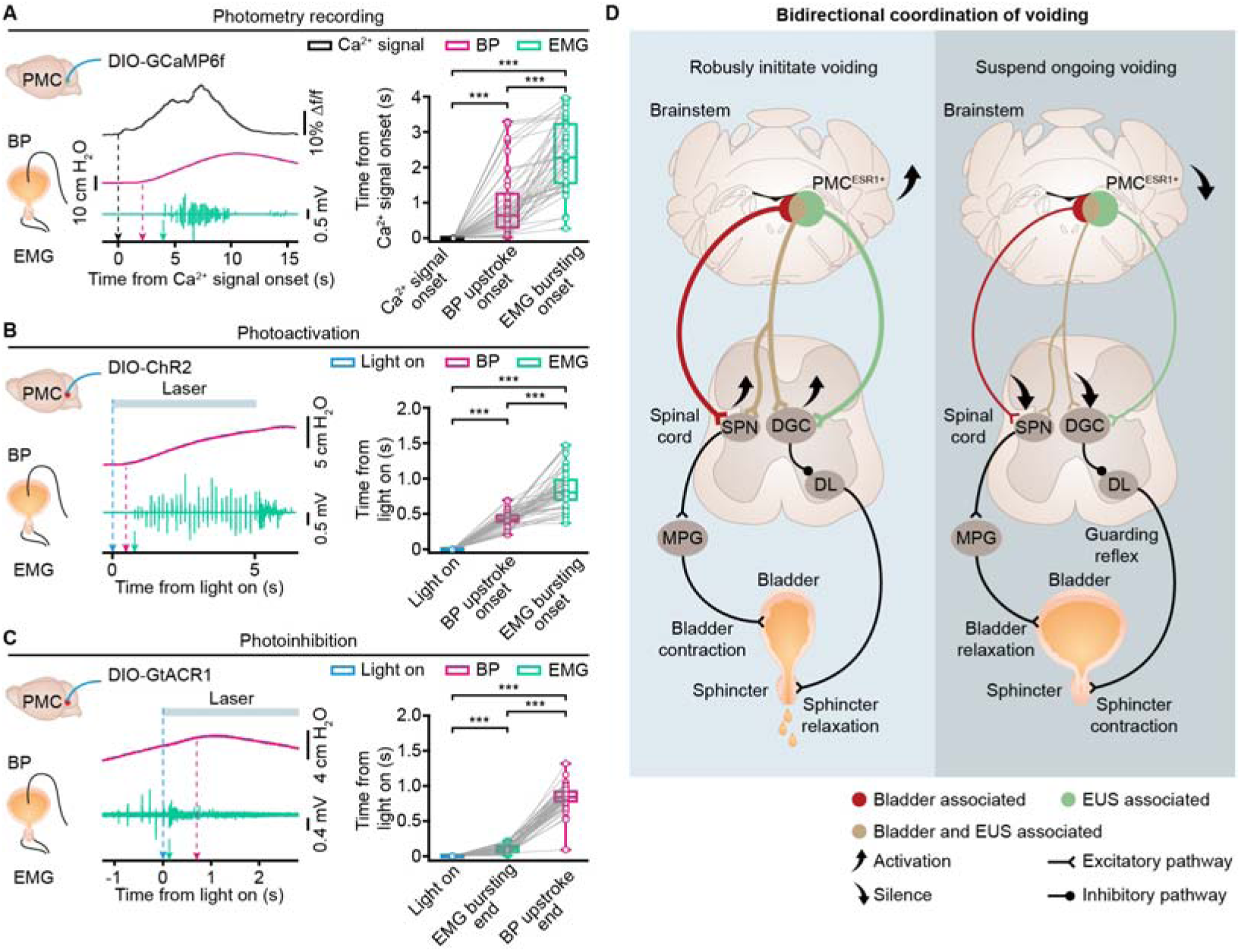
Coordination of bladder contraction and sphincter relaxation for urination by PMC^ESR1+^ cells. (A) Left: Example (left, with arrows indicating the onset timepoints) and quantification (right) of the temporal relationships among the onset times of Ca^2+^ signals, bladder pressure (BP) upstroke, and EUS-EMG bursting in photometry recordings (n = 46 trials from 8 mice; Ca^2+^signals onset, 0\0-0 s; BP upstroke onset, 0.6\0.3-1.2 s; EUS-EMG bursting onset, 2.3\1.5-3.2 s; \*\*\**P* = 3.5e-9 for all, Wilcoxon signed-rank test). (B) Example (left, with arrows indicating the onset timepoints) and quantification (right) of the temporal relationships among the onset times of light stimulation, bladder pressure upstroke, and EUS-EMG bursting for the photoactivation group (n = 50 trials from 10 mice; Light on, 0\0-0 s; BP upstroke onset, 0.4\0.4-0.5 s; EUS-EMG bursting onset, 0.8\0.7-1.0 s; \*\*\**P* = 7.6e-10 for all, Wilcoxon signed-rank test). (C) Example (left, with arrows indicating the onset timepoints) and quantification (right) of the temporal relationships among the onset time of light, EUS-EMG bursting end, and bladder pressure upstroke end for the photoinhibition group (n = 44 trials from 8 mice; Light on, 0\0-0 s; EUS-EMG bursting end, 0.1\0.1-0.2 s; BP upstroke end, 0.8\0.8-0.9 s; \*\*\**P* = 7.6e-9 for all, Wilcoxon signed-rank test). (D) A working model of the brainstem-spinal compound circuit for bidirectional and coordinated control of voiding. Abbreviations: SPN, sacral parasympathetic nucleus; DGC, dorsal gray commissure; DL, dorsolateral nucleus; MPG, major pelvic ganglia. The guarding reflex is triggered by bladder afferents via the pelvic nerves and mediated by spinal interneuronal circuits, which activate the urethral sphincter to prevent involuntary bladder emptying (Fowler et al., 2008). For all data points in (A, B), and (C), whisker-box plots indicate the median with the 25%-75% percentile as the box, and whiskers represent the minimum and maximum values.

Intriguingly, in photoinhibition experiments, the EUS-EMG bursting activity ended almost instantaneously, whereas the bladder pressure upstroke terminated slightly later (EMG bursting end: 0.1\0.1-0.2 s; bladder pressure upstroke end: 0.8\0.8-0.9 s, n = 44 events pooled from 8 mice; Figure 8C). This delay likely arises because the residual urine stream continues to excite urethral afferents, sustaining detrusor contraction via the urethra–detrusor facilitative reflex and thus prolonging the decline in bladder pressure (Barrington, 1921; Sasaki, 2004). Furthermore, factors such as measurement-related delays in bladder-pressure recording or lower spinal reflex pathways linking bladder afferents to sphincter control could influence the relative timing between bladder and sphincter activity (Chang et al., 2007; de Groat et al., 2015). Therefore, PMC^ESR1+^ cells ensure efficient urination by coordinately controlling bladder contraction and sphincter relaxation. Specifically, activation of PMC^ESR1+^ cells initiate bladder contraction followed by sphincter relaxation, enabling efficient urine elimination, whereas silencing these neurons after voiding onset causes the sphincter to close first and bladder contraction to cease, interrupting voiding (Figure 8D).

## Discussion

A prominent aspect of our data is that voiding events 100% correlated with PMC^ESR1+^ neuronal activity (Figure 1), and both photoactivation and photoinhibition of PMC^ESR1+^ cells yielded a 100% success rate in initiating and suspending the process of voiding, respectively (Figure 2 and Figure supplement 7). Whilst there are many other factors from both the lower spinal circuit and the higher cortical/subcortical circuits together to determine in what condition to void (de Groat, 2009; de Groat et al., 2015; Yao et al., 2018; Zderic, 2019; Mukhopadhyay & Stowers, 2020; Sartori et al., 2022), the instantaneous execution to efficiently initiate or suspend a voiding process involves a significant contribution of PMC^ESR1+^ cells in the brainstem as demonstrated here in this study. The bidirectional control, i.e., in the one way to initiate voiding whenever needed and suitable (which is a basic life need) (Mukhopadhyay & Stowers, 2020), and in the other way to suspend an ongoing voiding when needed (e.g., to release only a small volume of urine for landmarking) (Desjardins et al., 1973; Keller et al., 2018; Hyun et al., 2021), can be executed at 100% reliability by PMC^ESR1+^ cells given that all downstream nerves and muscles are intact and operational.

Classic studies have reported that lesions in the PMC or spinal cord produce urinary retention and detrusor-sphincter dyssynergia characterized by involuntary sphincter contractions during bladder contraction (Manente G, 1996; Bacsu et al., 2012; Panicker et al., 2015; Taweel & Seyam, 2015; Shimizu et al., 2023). These findings highlight the importance of PMC spinal-projecting neurons and their axons in coordinated motor control of the bladder and the EUS. In the present study, multiple lines of functional evidence deepen this concept by demonstrating that PMC^ESR1+^ neurons command both the bladder and the EUS. First, robust increases in PMC^ESR1+^ neuron activity are tightly correlated with bladder contraction and EUS relaxation during reflexive urination (Figure 1). Second, optogenetic activation of these neurons evokes both bladder contraction and EUS relaxation when the bladder is not empty, as reflected by elevated intravesical pressure and the emergence of EUS-EMG bursting (Figure supplement 7). Third, photoactivation of PMC^ESR1+^ neurons fails to elicit bladder contractions after pelvic nerve transection, yet still produces them following pudendal nerve transection (Figures 3-4). Fourth, voiding efficiency of the urination evoked by activation of PMC^ESR1+^ neurons is significantly reduced in pelvic- or pudendal-transected mice compared with sham controls, underscoring the necessity of coordinated bladder-sphincter action for efficient urine expulsion (Figures 5-6). Collectively, these results identify PMC^ESR1+^ neurons as a dedicated brain-output population that transforms integrated pro-urination commands into the coordinated contraction of the bladder detrusor and relaxation of the EUS required for normal voiding.

Furthermore, based on population dynamics obtained by fiber photometry (Figures 1D-1H, Figure supplements 1A-1F, and Figure supplements 11A-11H) and single-neuron firing properties recorded via optrode (Figures 1A-1C), we propose several mechanistic models for the engagement of dual- and single-projecting PMC^ESR1+^ neurons during natural micturition. One possibility is that all three populations (dual-projecting, SPN-projecting and DGC-projecting neurons) are co-activated, with the dual-projecting subset acting as a “bridging amplifier” that sustains rising bladder pressure while coordinating EUS relaxation. Alternatively, SPN-projecting neurons may be recruited first to initiate bladder contraction, followed by DGC-projecting neurons that evoke EUS bursting and facilitate urine entry into the urethra; once flow begins, the urethro-detrusor facilitative reflex could recruit dual-projecting neurons to further enhance voiding efficiency. In addition, contextual or state-dependent urination—such as scent-marking behavior characterized by multiple voiding events with smaller volumes than reflexive urination (Kaur et al., 2014; Malykhina, 2017; Mukhopadhyay & Stowers, 2020)—may predominantly rely on sequential and cooperative activation of single-projecting neurons. Other recruitment sequences remain conceivable. Future studies combining diverse urination-related behavioral paradigms with simultaneous recordings from projection-specifically labeled PMC neurons will be required to validate and refine these models.

Although PMC^ESR1+^ cells are capable of executing dynamic, real-time control of voiding with complete reliability, this does not imply that other cell types in the PMC are dispensable for perfect urination control. Rather, we propose that proper baseline control of bladder pressure is also essential, such as by CRH+ cells, which constitute the majority of PMC neurons and primarily regulate the bladder via the pelvic nerve while being modulated by various contextual factors (Vincent & Satoh, 1984; Wood et al., 2009; Hou et al., 2016). After all, it is not the marker gene ESR1 itself, which is abundantly expressed in many other brain regions and involved in diverse physiological and cognitive functions (Fang et al., 2018; Karigo et al., 2021; Liu et al., 2022), but rather the specific innervation pattern (Figure 8D) that enables this role in urination coordination. Specifically, any cell located within this brainstem nucleus (PMC) (Kawatani et al., 2021) that possesses dual innervations targeting both the bladder and sphincter can serve as a potent contributor to urination coordination (Griffiths, 2015), irrespective of its molecular identity.

Our optrode data (Figure 1) show that, at the single-cell level, less than half (11/28) of ESR1^+^ cells exhibit strong correlations with voiding events, although this proportion is still much higher than that of non-ESR1^+^ cells (9/51). These observations suggest that the functional recruitment of PMC neurons during urination is determined not merely by molecular identity (ESR1 expression), but rather by their specific projection patterns and circuit connectivity. However, testing this hypothesis is currently hindered by technical constraints. We are unable to perform projection-specific, single-cell-identified functional profiling across the ESR1+ cells or the entire PMC population. Specifically, selective labeling and classification of PMC neurons based on their distinct spinal projection targets to correlate anatomical connectivity with physiological response dynamics remains to be achieved. Additionally, our study is limited by the methodology used to establish temporal sequences, which relies on simultaneous recordings of bladder pressure and EUS-EMG. While EUS-EMG provides direct and rapid readout of sphincter activity, bladder pressure measurements may not precisely reflect pelvic-nerve-driven contractions due to mechanical delays in pressure transmission. Future studies incorporating direct pelvic nerve recordings alongside EUS-EMG monitoring will be essential to validate the precise timing of bladder-sphincter coordination.

## Conclusions

In summary, we have identified the essential role of PMC^ESR1^ neurons in coordinating urination through co-innervation of both the bladder and urethral sphincter. Our findings offer new insights into the anatomical and physiological basis and research paradigms for the coordinated engagement of parasympathetic and somatic functions of urination control. PMC^ESR1^ neurons may serve as a key focal point for advancing our understanding of the neural mechanisms underlying urination, both in physiological contexts and pathological conditions such as brain injuries, spinal cord injuries, and peripheral nerve damages.

## Materials and Methods

### Animals

The experiment procedures were approved by the Third Military Medical University Animal Care and Use Committee and were conducted strictly in adherence to established guidelines. This study utilized Wild-type C57BL/6J mice, ESR1-IRES-Cre (Jackson Laboratory, stock #017911) (Lee et al., 2014) and CRH-IRES-Cre (Jackson Laboratory, stock #012704) (Chen et al., 2015) mice. These mice were group-housed in an environment-controlled room at 23-25℃ and 50% humidity, with 4-5 mice per cage, on a 12-hour light/dark cycle, and with free access to food and water. Mice implanted with optical fibers were housed individually. Both male and female mice, aged 8-20 weeks, were randomly assigned to various experiments. The figure legends detail the number of animals utilized in each experiment.

### Virus vectors and CTB

The study utilized the following viruses and cholera toxin subunit B (CTB) for various experiments: For fiber photometry recording experiments, AAV2/9-DIO-GCaMP6f (titer: 0.5 × 10^12^ vg/ml) was unilaterally injected into the PMC. For optogenetic manipulation experiments, AAV2/9-hsyn-DIO-hGtACR1-mCherry (titer: 1.43 × 10^13^ vg/ml) and AAV2/8-DIO-ChR2-mCherry (titer: 1.33 × 10^13^ vg/ml) were bilaterally delivered into the PMC for photoinhibition and photoactivation, respectively. For retrograde tracing experiments, pAAV2/retro-EF1a-DIO-EGFP (titer: ≥ 1.00 × 10^12^ vg/ml) and pAAV2/retro-EF1a-DIO-mCherry (titer: ≥ 1.00 × 10^12^ vg/ml) were injected into the spinal cords of both ESR1-Cre and CRH-Cre mice. To better visualize the injection sites, CTB-647 (0.2%, C34778, Thermo Fisher) was co-injected with the virus at a 1:10 volume ratio in some experiments. For anterograde tracing experiments, AAV2/9-hEF1a-fDIO-mGFP (titer: 1.00 × 10^13^ vg/ml) was injected unilaterally into the PMC, and a 10: 1 volume mixture of AAV2/2Retro-hsyn-FLEX-Flpo (titer: 1.00 × 10^13^ vg/ml) and CTB 555 (0.2%, C34776, Thermo Fisher; used solely for determining the injection site) was injected into the spinal cord. For photometry recording of dual-projecting neurons, rAAV/9Retro-hSyn-SV40 NLS-Cre (titer: 5.01 × 10^12^ vg/ml) and AAV2/2Retro-hSyn-FLEX-GCaMP6s (titer: 1.73 × 10^13^ vg/ml), each mixed with CTB-555 (0.2%, C34776, Thermo Fisher) at a 10:1 volume ratio, were delivered into the SPN and DGC in the lumbosacral spinal cord, respectively. For photoactivation of dual-projecting neurons, rAAV/9Retro-hSyn-SV40 NLS-Cre (titer: 5.01 × 10^12^ vg/ml) and AAV2/2Retro-hEF1a-DIO-hChR2-mCherry (titer: > 1.00 × 10^13^ vg/ml), each mixed with CTB-488 (0.2%, C34775, Thermo Fisher) at a 10:1 volume ratio, were delivered into the SPN and DGC in the lumbosacral spinal cord, respectively. In control experiments, rAAV2/9-EF1a-DIO-EYFP (titer: 4.20 × 10^12^ vg/ml) and rAAV2/9-EF1a-DIO-mCherry (titer: 5.28 × 10^12^ vg/ml) were used. All viruses were purchased from Obio Biotechnology Co., Ltd. (Shanghai, China), Taitool Bioscience Co., Ltd. (Shanghai, China), BrainVTA Co., Ltd. (Wuhan, China), or Braincase Co., Ltd. (Shenzhen, China).

### Stereotaxic injections and optical fiber implant

To target PMC^ESR1+^ neurons, ESR1-Cre mice were anesthetized with isoflurane (3% for induction and 1.5–2% for maintenance) and head-fixed in a stereotaxic frame (RWD Life Science Co., Ltd.; Shenzhen, China). Body temperature was maintained at 36°C throughout the surgery using a heating pad. Local anesthesia (lidocaine, 6 mg/kg, subcutaneous injection) was administered at the incision site before making the incision. A small cranial hole above the PMC was created using a dental drill. Approximately 80 nl of the viral solution was delivered to the injection site at a controlled rate of 20 nl/min, either unilaterally or bilaterally, using a micro-syringe pump connected to a glass pipette. The coordinates for PMC injections were: anteroposterior (AP) is −5.45 mm, mediolateral (ML) is ± 0.70 mm, and dorsoventral (DV) is −3.14 mm from the dura. The pipette was then slowly withdrawn over 5 min to prevent virus overflow. The incision was closed with sutures, and the mice received antibiotics and analgesics post-surgery. A heating pad was used for the mice to aid recovery from anesthesia. The mice were then group-housed in their home cages for 3-4 weeks to allow for viral expression.

For optical fiber implantation, the optical fiber (NA: 0.48, diameter: 200 μm, Doric lenses, Quebec City, QC, Canada) was fixed into a metal cannula and positioned 50 μm above the PMC injection site, with coordinates of AP, −5.45 mm; ML, ± 0.7 mm; and DV, −2.95 mm from the dura mater (Figures 1-4 and Figure supplements 1, 2, 6, 8, 9, and 11). Notably, in some experiments, the left optical fiber was implanted at a 33° lateral angle targeting the PMC, with coordinates of AP, −5.45 mm; ML, +2.5 mm; and DV, −3.0 mm from the dura mater (Figure supplement 7). The fiber was implanted unilaterally for photometry experiments and bilaterally for optogenetics experiments. Dental cement was used to affix the optical fiber to the skull. Mice were housed individually and given 3-5 days to recover before recording or stimulation sessions. After the experiments, the placement of the virus and optical fiber was confirmed by histology in each mouse.

### Targeted spinal cord injections

For spinal cord injection surgery, the method previously described was used (Chen et al., 2019). Briefly, ESR1-Cre, CRH-Cre, and C57BL/6J mice were anesthetized under isoflurane (3% for induction and 1.5–2% for maintenance) and positioned on a heating pad. After shaving the hair, a midline skin incision (1-2 cm) was performed over the lumbar segments following local anesthesia (lidocaine, 6 mg/kg, subcutaneous injection) at the incision site. The tissue and muscle connected to the dorsal spine were dissected to expose the T12-L2 vertebrae. The spine was affixed to a stereotactic frame using a spinal adapter (68094, RWD), and the spinal cord between L1 and L2 was exposed by removing the ligamentous and epidural membranes. Using the central vein as a reference, a total of 80 nL of virus solution was injected at two different locations, each injection (40 nL) was administered through a micro-syringe pump connected to a glass pipette, targeting DGC (+ 0.12 mm lateral to the central vein, 0.52 mm deep from the dorsal surface at a 10° lateral angle) and SPNs (−0.12 mm lateral to the central vein, 0.56 mm deep from the dorsal surface at a 30° lateral angle). The pipette remained in position for a minimum of 5 min before being slowly removed to prevent any leakage. The injection sites were sealed with tissue glue (Vetbond, 3M Animal Care Products), followed by suturing of the skin. Mice received antibiotics and analgesics post-surgery. Approximately 3-4 weeks after the injections, the brain and spinal cords were extracted for histological validation. In ESR1-Cre or CRH-Cre mice, PMC cells projecting to the SPN (labeled with mCherry) and DGC (labeled with EGFP) regions of the spinal cord were manually quantified using Image J.

### Pudendal nerve and pelvic nerve transection

Pudendal nerve transection was performed as described previously (Khorramirouz et al., 2016; Peh et al., 2018). Briefly, mice were anesthetized with 1.5-2% isoflurane in oxygen and positioned on a heating pad. A midline skin incision was made along the back from L4 to the coccyx, followed by paraspinal incisions through the gluteal muscles and fascia to expose the sciatic nerve. The sciatic nerve was gently retracted to expose the pudendal nerve. Using microsurgical scissors, the bilateral pudendal nerves, along with the anastomotic branch, were carefully dissected and excised.

For pelvic nerve transection, modifications to the established method (Chang et al., 2018) were made to minimize surgical trauma. Mice were positioned laterally, and bilateral paraspinal incisions were extended upward. The sciatic and pudendal nerves were gently retracted laterally to expose the pelvic nerve, which was identified (originating from the sacral segments of the spinal cord and connecting to the major pelvic ganglion) and severed. For experiments in freely moving mice, following pelvic or pudendal nerve transection, the muscle and skin layers were sutured, and postoperative antibiotics and analgesics were administered. In the sham groups, the same procedures were performed except that the pudendal or pelvic nerves were left intact.

### Fiber photometry recording and analysis in freely behaving mice

The Ca^2+^ recordings of PMC^ESR1+^ neurons were conducted using a fiber photometry setup, as described previously (Yao et al., 2018; Rao et al., 2022). Fluorescence at the fiber tip was excited by blue light (0.22 mW/mm^2^). Mice with implanted fibers were injected intraperitoneally with diuretics (furosemide, 40 mg/kg) and acclimated in a testing chamber equipped with a bottom camera (1,280 × 720 pixels) for 20 min before recording. Signals from PMC^ESR1+^ neurons and voiding behavior were simultaneously recorded for approximately 40 min. Ca^2+^ signals were sampled at 2000 Hz using NI LabVIEW software (National Instruments, USA), while behavioral video was captured at 30 Hz. Fiber photometry data and video were synchronized via event markers. For data analysis, all signals were processed with a Savitzky-Golay filter (third-order polynomial, 50 side points) for low-pass filtering. Δf/f = (f - f_baseline_)/f_baseline_ was calculated to assess photometry signals during voiding, where f_baseline_ represents the minimum fluorescence recorded. Results were presented as heatmaps using MATLAB. Ca^2+^ signal data were shuffled by dividing the original dataset into 10 segments and randomly associating them with voiding events. Positive signals were defined as Ca^2+^ signal amplitudes exceeding three times the noise band (the standard deviation). This procedure was also applied to control mice to correct for movement artifacts.

### Single-unit with optrode recording and analysis

To identify the single-unit activity of PMC^ESR1+^ neurons, optrode recordings were performed as described previously (Qin et al., 2018; Qin et al., 2022; Yang et al., 2023). Briefly, the optrode consisted of a 200 µm optical fiber and four tetrode assemblies aligned in a line, spaced 100 µm apart. The optical fiber was secured to the electrodes, positioned 500 µm above their tips, and connected to an LED. Each electrode assembly comprised four twisted tungsten wires (25 µm, California Fine Wire), allowing vertical movement via micromanipulators. Optrode implantation surgery was performed in ESR1-Cre mice expressing ChR2 in the PMC, with the electrode tips aligned and implanted 2.80 mm below the brain dura. After a recovery period of 5-7 days, the tetrodes were slowly inserted to a target depth of −2.95 mm and recording began. Single-unit signals from PMC^ESR1+^ neurons in freely behaving mice were recorded using an RHD2000 USB board (C3100, Intan Technology) at 20 kHz, while behavioral video was captured simultaneously. Units with short spike latencies (< 7 ms) in response to light pulses of varying intensities (5 mW, 10 mW, 15 mW, and 20 mW) and high responsiveness (> 70 %) were identified as PMC^ESR1+^ cells. To confirm recording locations, electro-lesions were performed by applying a current (10 μA, 12 s) through the tetrodes.

The raw recorded data were preprocessed using established methods to extract peaks (Qin et al., 2018). All events exceeding the amplitude threshold (set at four standard deviations above the background) were kept for further analysis. The average firing rate of each cell was calculated within a sliding time bin of 20 seconds around voiding (0.1 s intervals), divided by the total number of trails, and adjusted by subtracting the baseline value (the median firing rate during the −10 to −5 s before voiding). The results were visualized as a heatmap (logarithmic analysis) in Figure 1. Statistical analysis of the average firing rates over a 2-second interval before voiding (from −10 s to −8 s) and around voiding (from −1 s to 1 s) was conducted for PMC^ESR1+^ neurons and non-PMC^ESR1+^ neurons, respectively.

### Optogenetic experiments in freely behaving mice

Before optogenetic stimulation, mice underwent the same procedures as for fiber photometry: they received a diuretic with the intraperitoneal injection to increase urination events and were acclimated to the testing chamber for at least 20 min. For optogenetic inhibition experiments, bilateral stimulation (1 mW/mm² at the fiber tips) was delivered using 473 nm blue light. To assess the effect of light inhibition on urination, mice were placed in a glass chamber (28 cm × 16 cm × 30 cm) with a 0.19 mm filter paper (14.6 cm × 27 cm, BWD) underneath. Urination was observed at three stages: pre-photoinhibition (light-off), during the 5 s or 60 s of constant photoinhibition (light-on), and post-photoinhibition (light-off). The photoinhibition parameters (frequency and duration) were controlled via the NI LabVIEW platform (National Instruments, USA). Real-time urination behavior was monitored simultaneously using two cameras positioned above and below the glass chamber. Closed-loop photo-inhibition was manually triggered upon the first visible urine patch detected via real-time video. Trials with triggers >2.51s after urination onset were excluded. Urination cessation was defined as the initiation of movement following urination. The void area, total void duration, and latency were analyzed from the video and are presented in Figure 2 and Figure supplements 5-6.

For optogenetic activation experiments, bilaterally fiber-implanted mice were connected to two 473-nm blue laser generators (5 mW/mm^2^ at fiber tips, 25 Hz frequency, 15 ms pulses) via optic fibers. During testing, blue light was delivered for 5 s with approximately 3-min intervals between trials, over a 30-45 min session. This light stimulation was repeated over two days with a one-day interval. Mouse behavior was monitored and recorded simultaneously. The stimulation was performed before and after transection for photoactivation experiments in freely behaving mice that underwent pelvic or pudendal nerve transection, following the same procedures. Note that the photostimulation experiments for pelvic or pudendal nerve transection were conducted one day after the surgery. Detailed success rates, latency of voiding, and void area after photoactivation are shown in Figures 5-6 and Figure supplement 7.

### Simultaneous cystometry and electromyography in freely behaving and anesthetized mice

Bladder catheter and urethral sphincter electrode implantation were performed as previously described (Hou et al., 2016; Keller et al., 2018; Verstegen et al., 2019). Briefly, adult fiber-implanted ESR1-Cre mice were anesthetized with isoflurane and placed on a heating pad. A lower mid-abdomen incision exposed the bladder and urethral sphincter. PE-10 tubing was inserted through the bladder dome and secured with a 6-0 Ethicon suture. For electromyography (EMG) recording, two 160 μm silver-plated copper wire electrodes (P/N B34-1000, USA) with 2 mm exposed tips were inserted into the external urethral sphincter (EUS) on both sides using a 30-gauge needle, positioned between the urethra and pubic symphysis, and spaced at least 2 mm apart. A ground wire with 4 mm exposed tips was inserted subcutaneously near the sternal notch to minimize signal interference. The ends of the bladder catheter tubing and wire electrodes were exteriorized through an incision on the skin at the back of the neck, and both abdominal and neck incisions were closed. For measuring intravesical pressure, the bladder tubing was connected to a pressure transducer (YPJ01H; Chengdu Instrument Factory, China) and a syringe pump (RWD404; RWD Technology Corp., Ltd., China) via three-way stopcocks. Bladder pressure and EUS-EMG data were recorded through a multi-channel physiological recording device (RM6240; Chengdu Instrument Factory, China) sampled at 8 kHz. After surgery, mice were permitted to recover from anesthesia and resume walking.

For simultaneous recording, all fiber-implanted mice were anesthetized with urethane (1.2 g/kg, i.p.) and continuously infused with room-temperature physiological saline at 30-50 µl/min via the bladder catheter for at least 45 min. Recording or stimulation was performed once regular bladder pressure cycles associated with natural urination events were established. “Filling bladder” was defined as continuous saline infusion, while “non-filling bladder” was defined as no infusion. Cystometry and EUS-EMG data were captured via commercial acquisition software, alongside monitoring of Ca^2+^ signals and mouse behavior. For cross-correlation analysis, cystometric data, EMG data, and photometry data were first downsampled to 40 Hz and standardized using z-scored. The original EMG data were processed to extract their envelope using MATLAB’s “envelope” function. The original data were merged, and divided into 10 segments, and segments of photometry data were randomly matched with segments of cystometry or envelope EMG data to create shuffled datasets. Cross-correlations of cystometric and photometry data, or EMG and photometry data were calculated using MATLAB’s “xcorr” function, and the peak values of cross-corrections were reported, as shown in Figure 1. Photometry recording from SPN- and DGC-projecting PMC neurons was performed during simultaneous cystometry and EUS-EMG, following the procedures and analysis described above. Results are shown in Figure supplement 11.

For photoactivation experiments performed simultaneously with cystometry and electromyography recording, blue light pulses were delivered periodically (every 30 s) at 25 Hz for 15 ms, lasting 5 s, or randomly at intervals between 20 s and 40 s, with each condition repeated at least 15 times. These parameters were chosen to ensure reliable spiking of PMC^ESR1+^ neurons without inducing depolarization block and to effectively drive urination (Keller et al., 2018). For photoactivation experiments performed simultaneously with cystometry and urethral electromyography recording under pelvic or pudendal nerve transection conditions, the surgical procedures and pre-recording preparations were the same as those described above, with the following changes: Mice underwent three stages of randomized photostimulation (consisted of 25 Hz, 15 ms, 5-s durations, with intervals between 30 s and 60 s) in sequence under urethane anesthesia (1.2 g/kg, i.p.): an intact nerve period, a pudendal nerve or pelvic nerve transection period, and a period with both pudendal nerve and pelvic nerve transection. All trials were pooled to assess the impact of nerve transection on bladder and sphincter function for each mouse. For photoactivation of SPN- and DGC-projecting PMC neurons in freely moving mice, previously described procedures were used with minor modifications. Mice were placed in a glass chamber (28 cm × 16 cm × 30 cm) lined with 0.19 mm filter paper and voiding events were monitored in real time using two cameras positioned above and below the chamber. Cystometric parameters were analyzed using MATLAB. For photoactivation, ΔP = P_5 sec_ - P_0 sec_ (where P_0 sec_ is the pressure at the onset of laser stimulation and P_5 sec_ is the pressure at the end of laser stimulation) was calculated to assess the immediate effect of light activation on bladder pressure, and the pressure ratio P_ratio_ = P_max (0-5)sec_ / P_mean (−5-0)sec_ (where P_max (0-5)sec_ is the maximum pressure from laser onset to cessation, and P_mean (−5-0)sec_ is the average pressure during the 5 seconds preceding laser onset) was used to quantify the relative pressure change. The EUS-EMG data (burst duration and area under the curve) were analyzed using multi-channel physiological recording software. The spectrogram was generated using envelope EMG data around photostimulation in MATLAB, as shown in Figures 3-4, and Figure supplement 9. The onset time of bladder pressure upstroke was identified by finding the maximum of the second derivative of cystometry curves around pressure peaks using a MATLAB script. The onset time of EMG bursting (Cheng & de Groat, 2004; Kadekawa et al., 2016) was manually defined as the points at which bursting activity begins. The results are shown in Figure 8 and Figure supplement 12.

For photoinhibition experiments performed simultaneously with cystometry and electromyography recording, the light was manually triggered at the onset of EMG bursts and delivered as 5 s or 60 s of constant photoinhibition to ensure continuous neuronal suppression. Cystometric parameters, including Δpressure = P_peak_ – P_min_ (where P_peak_ is the peak pressure and P_min_ is the minimum pressure after the peak), were used to quantify the extent to which the ongoing pressure wave was aborted by photoinhibition. The threshold pressure (bladder pressure upstroke) was determined manually using multi-channel physiological recording software. The EUS-EMG data (burst duration, area under the curve, and latency of termination) was analyzed using the multi-channel physiological recording software. The latency of bursting termination is defined as the interval from laser onset to the end of EUS-EMG bursting. The end time of bladder pressure upstroke (termination of the rapid increase in bladder pressure before the end of voiding, denoting bladder relaxation) was identified by finding the minimum of the second derivative of cystometry curves around pressure peaks using a MATLAB script. The end time of EMG bursting was manually defined as the points at which the onset of tonic activity after bursting. The results are shown in Figure 8.

### Histology and immunohistochemistry

Following the completion of all experiments, histological verification of fiber implantation or virus injection positions was conducted. Mice were deeply anesthetized with 1% sodium pentobarbital (10 ml/kg), followed by transcardial perfusion with cold 0.9% saline and 4% paraformaldehyde (PFA). The brain or spinal cord (in some experiments) was then extracted and post-fixed overnight at 4 °C in ice-cold 4% PFA. Coronal brain sections (40 µm) and the thoracolumbar and lumbosacral segments of spinal cord sections (70 µm) were cut using a freezing microtome. For TH immunohistochemistry, free-floating sections were initially incubated in 1% PBST (1% Triton X-100 in PBS, Sigma) for 60 min. The sections were blocked with 10% donkey serum (Sigma) in 0.1% PBST for 2 hours at room temperature. Following blocking, then incubated at 4°C for 24 hours with anti-TH antibodies (1:200 dilution, rabbit, Sigma-Aldrich). After extensive washing with PBS, sections were incubated with a secondary antibody (1:300, Alexa Fluor 488 or 594 donkey anti-rabbit, Invitrogen) at room temperature for 2 hours. Finally, all sections, including the target segments of the spinal cord, were incubated with DAPI (1:1000, Beyotime) for 15 min. Images were acquired using a confocal microscope (TCS SP5, Leica) equipped with × 10, and × 20 objectives, utilizing 405 nm, 488 nm, and 552 nm lasers, or using an Olympus microscope.

## Statistical analysis

All data were processed and statistically analyzed using Prism 8 GraphPad, MATLAB, and SPSS 22 software. For unpaired group comparisons, the Wilcoxon rank-sum test was used, and for paired groups, the Wilcoxon signed-rank test was applied, as described in the figure legends. Analysis was performed by investigators blinded to the experiments. The n value reflects the final number of animals in each experiment group, with animals excluded if histological verification of gene expression showed poor or absent. Data are represented as median with 25%-75% percentiles, and statistical significance was defined as * < 0.05, ** < 0.01, *** < 0.001; ns, no significant difference.

## Acknowledgments

The authors are grateful to Ms. Jia Lou for her help in composing and editing the layout of the figures. This study was supported by grants from the National Natural Science Foundation of China to X.C. (No. 31925018, 32127801), the National Key R&D Program of China to X.C. (2021YFA0805000), the Suzhou Science and Technology Plan Project (SZS2022008), and the Jiangsu Provincial Big Science Facility Initiative (BM2022010), and the Guangxi Talent Program (“Highland of Innovation Talents”). X.C. is a member of CAS Center for Excellence in Brain Science and Intelligence Technology.

## Additional information

### Funding

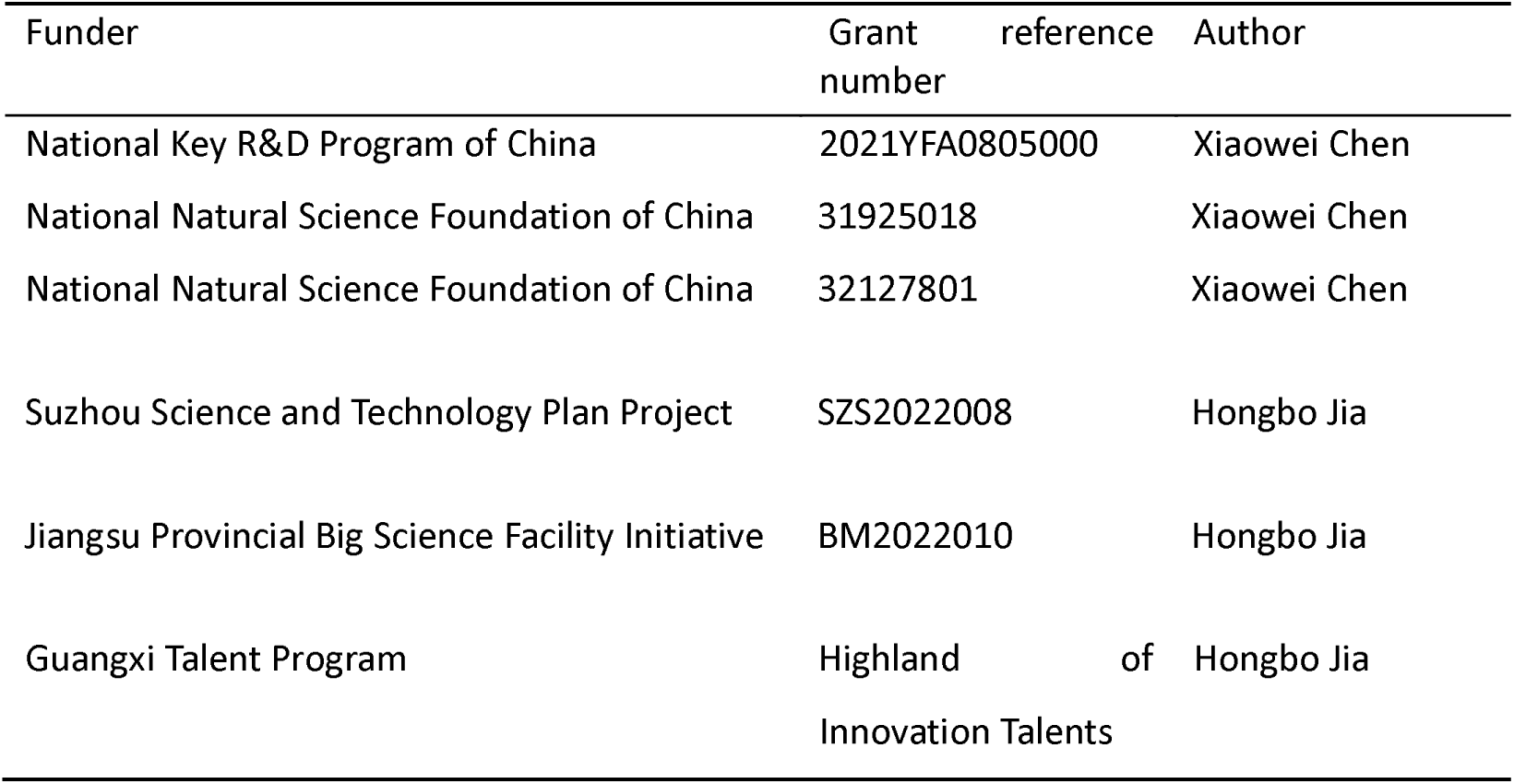

### Author contributions

Project design, J.W.Y, and X.C.; injection and histology, X.L., X.P.L., J.L.(1), L.X.Y., and J.L.(2); behavior experiments, X.L., C.H.Y., and X.W.; nerves transection, X.L.; fiber recording, X.L. C.H.Y., and X.P.L.; electrophysiology, X.L., H.Q., T.L.J., and X.W.; fiber recording, X.L. and L.X.Y.; cystometry and electromyography experiments, X.L., X.P.L., and J.L.(1); data interpretation and analysis, X.L., H.Q., S.S.L., H.B.J., X.Liao, J.W.Y., and X.C.; figure preparation, X.L., H.B.J., J.W.Y, and X.C.; manuscript writing, X.L., H.B.J., J.W.Y, and X.C. with the help of all co-authors. All authors read and commented on the manuscript.

### Ethics

All experiment procedures were approved by the Third Military Medical University Animal Care and Use Committee and were conducted strictly in adherence to established guidelines.

## Additional files

### Data availability

The supporting data underlying Figures 1-8 and Figure supplements 1-12 are provided as Source Data files. Any additional information underlying the findings of this study is available from the corresponding authors upon reasonable request.

### Code availability

The codes supporting the current study have not been deposited in a public repository, but are available from the corresponding author upon request.

## Figures supplement

**Figure supplement 1.**
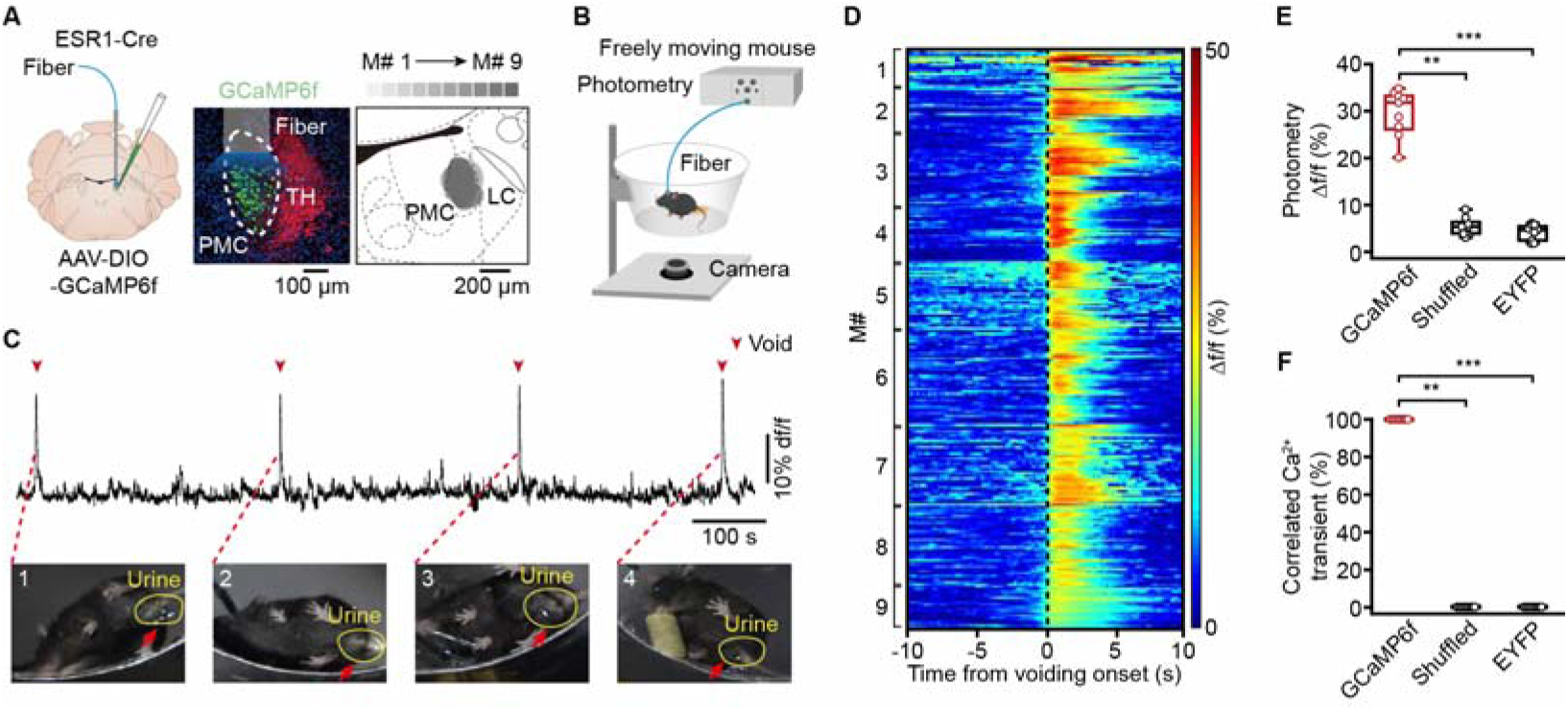
The activity of PMC neurons increases during successful voiding, related to Figure 1. (A) Schematic (left) of labeling and representative histology (middle) of PMC^ESR1+^ cells labeled with GCaMP6f. Scale bar: 100 µm. TH (tyrosine hydroxylase) stained neurons in the locus coeruleus. Right: Overlay of GCaMP6f-labelled areas from 9 mice. Scale bar: 200 µm. (B) Schematic of fiber photometry Ca^2+^ recording in a freely moving mouse. (C) Representative Ca^2+^ traces (top) and voiding events (bottom, yellow circle). (D) Cumulative sessions of Ca^2+^ signals aligned to voiding onset (dotted line). (E) Boxplots showing the amplitude of voiding-related Ca^2+^ signals in various groups (n = 9 mice per group, 31.8%\26.7%-33.2% for Gcamp6f, 5.4%\3.7%-6.0% for Shuffled, 4.7%\2.5%-5.7% for EYFP, \*\**P* = 3.9e-3 (Gcamp6f versus Shuffled), Wilcoxon signed-rank test; \*\*\**P* = 4.1e-5 (Gcamp6f versus EYFP), Wilcoxon rank-sum test). (F) Detected voiding-related Ca^2+^ events in various groups (n = 9 mice per group, 100%\100%-100% for Gcamp6f, 0%\0%-0% for Shuffled, 0%\0%-0% for EYFP. \*\**P* = 3.9e-3 (Gcamp6f versus Shuffled), Wilcoxon signed-rank test; \*\*\**P* = 4.1e-5 (Gcamp6f versus EYFP), Wilcoxon rank-sum test). For all data points in (E, F), whisker-box plots indicate the median with the 25%-75% percentile as the box, and whiskers represent the minimum and maximum values.

**Figure supplement 2.**
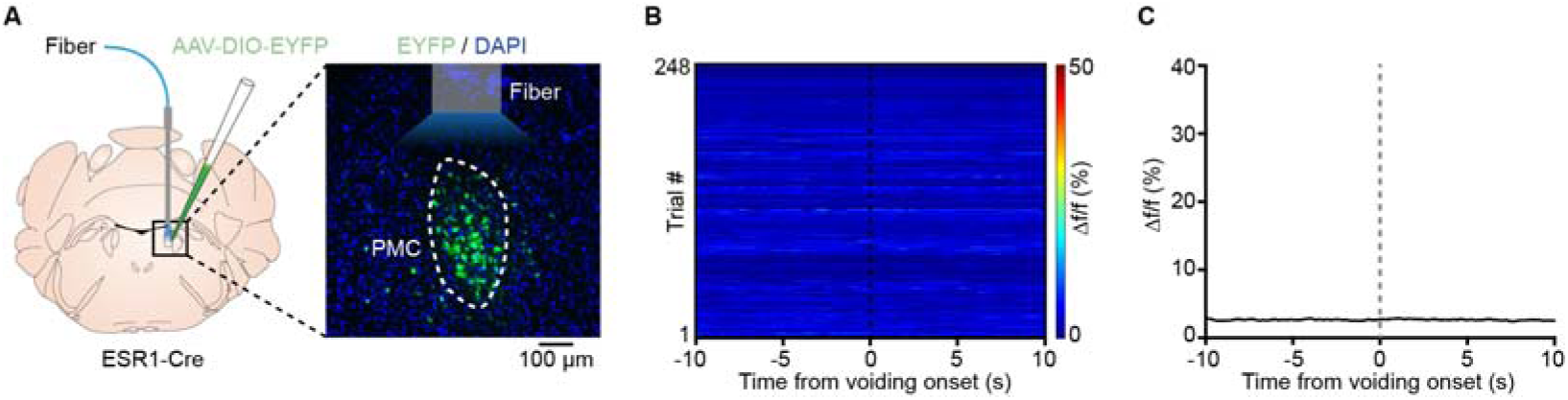
No change in fluorescence responses of EYFP-labeled PMC cells during voiding. (A) Schematics of labeling (left) and representative histology (right) of PMC^ESR1+^ cells labeled with EYFP. Scale bar, 100 µm. (B) Cumulative sessions of fluorescence responses aligned to voiding onset (n = 248 trials from 9 mice). (C) Average fluorescence traces of PMC^ESR1-EYFP^ aligned to voiding onset. The black line and shading represent mean ± s.e.m., respectively.

**Figure supplement 3.**
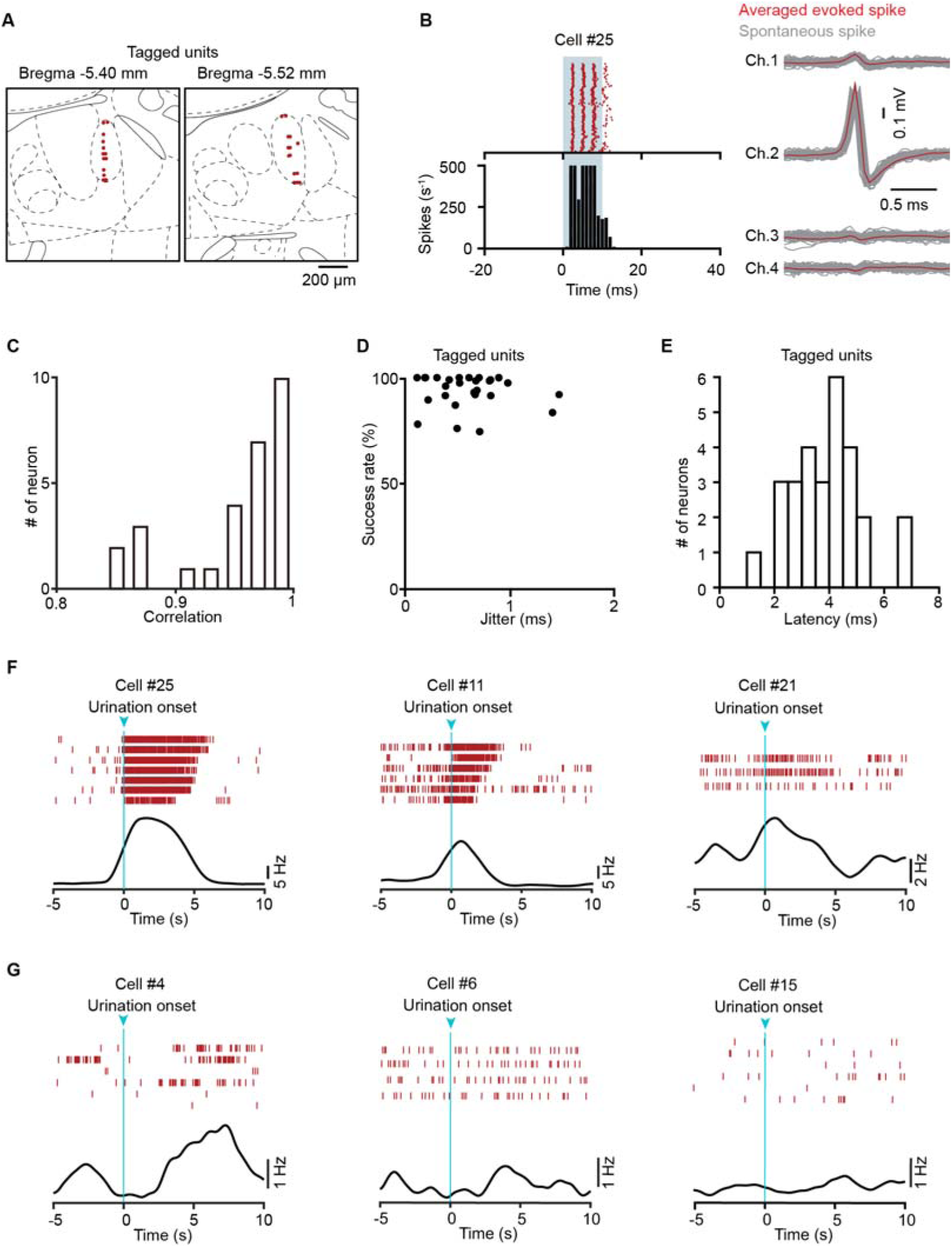
Identification of recorded PMC^ESR1+^ cells in optrode recordings. (A) Tetrode locations for all recorded PMC^ESR1+^ units (red dots, n = 28 units from 4 mice). (B) Left: Raster plots (top) and histograms (bottom) showing the firing patterns of representative PMC^ESR1-ChR2^ units upon optical stimulation at a power of 10 mW. Right: Waveforms of light-induced (red) and spontaneous (gray) spikes from the unit shown on the left. (C) Distribution of correlation coefficients between spontaneous and light-induced spikes across all recorded PMC^ESR1+^ units. (D) Comparison of success rates and temporal jitter for the first light-induced spike in all recorded PMC^ESR1+^ units. (E) Latency distribution for all recorded PMC^ESR1+^ units. (F, G) Representative firing patterns of opto-tagged PMC^ESR1+^ cells. F: Increased activity in all trials (left, middle) and in a subset of trials (right) during urination; G: No increase in activity during urination. Top: Raster plots. Bottom: Average firing rates.

**Figure supplement 4.**
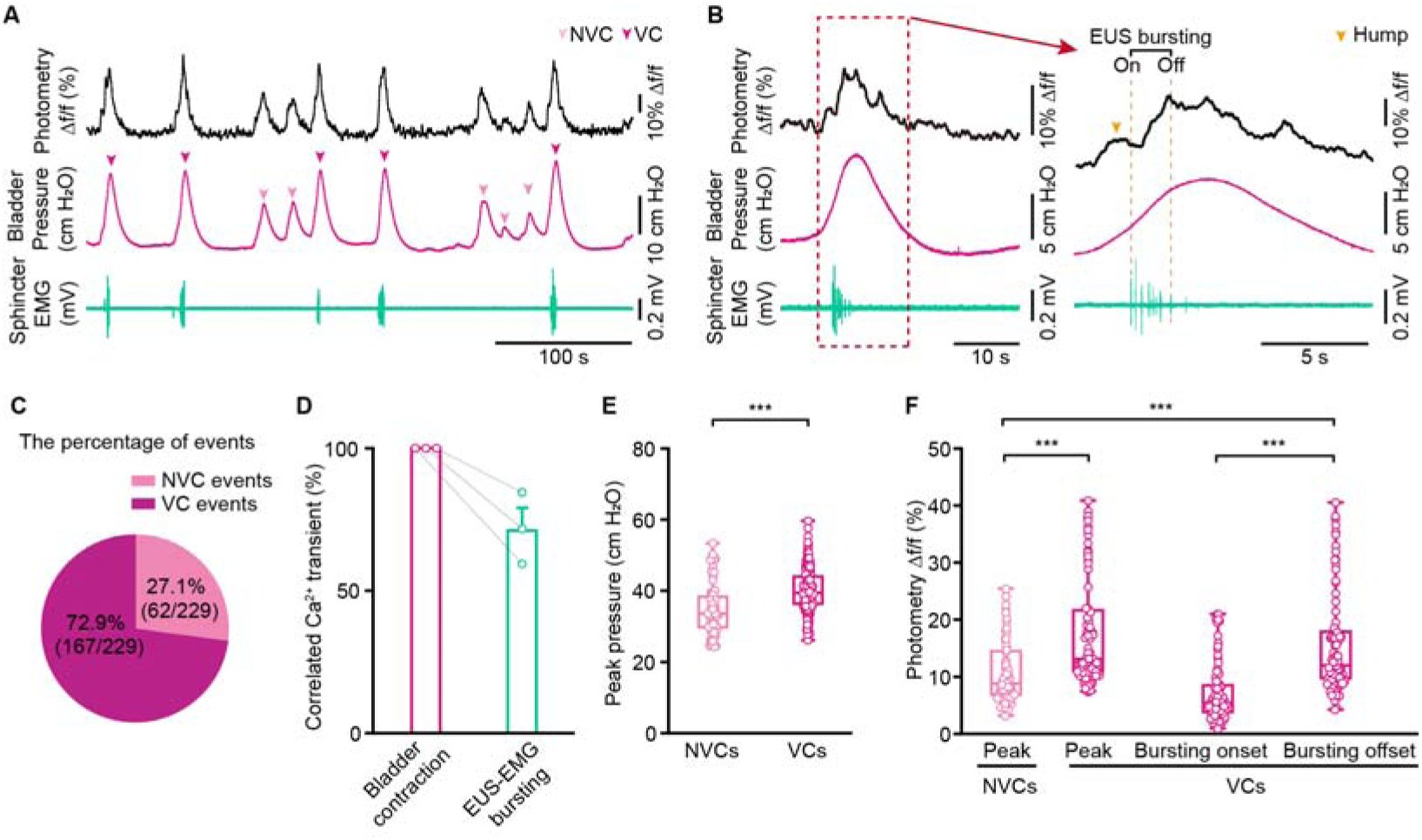
No voiding contractions (NVCs) correlate with Ca signals of PMC cells. (A) Representative traces of Ca^2+^ transients (black), bladder pressure (magenta), and EUS-EMG (teal) for no voiding contractions (NVCs) and voiding contractions (VCs). (B) Left: Representative fiber-photometry trace aligned to an EUS bursting episode. Right: Expanded view of the boxed region showing a small Ca²⁺ signal hump before bursting onset (yellow arrow) and a continuous rise during the bursting. (C) The percentage of NVCs (n = 62 events from 3 mice) and VCs events (n = 167 events from 3 mice). (D) Correlation rate of Ca^2+^ transient with bladder contraction and EUS-EMG bursting events (n = 3 mice per group). (E) Comparison of peak bladder pressure between NVCs and VCs (NVCs: n = 62 events from 3 mice, VCs: n = 167 events from 3 mice; \*\*\**P* = 6.03e-7, Wilcoxon rank-sum test). (F) Quantification of peak Ca²⁺ signals during NVCs and VCs, and Ca²⁺ signal amplitudes at bursting onset and offset during VCs (NVCs: 62 events from 3 mice, VCs: 79 events from 3 mice; \*\*\**P* = 8.2e-7, NVC peak versus VC peak, and \*\*\**P* = 4.4e-4, NVC peak versus VC bursting offset, Wilcoxon rank-sum test; \*\*\**P* = 1.1e-14, VC bursting onset versus VC bursting offset, Wilcoxon signed-rank test). For all data points in (E, F), whisker-box plots indicate the median with the 25%-75% percentile as the box, and whiskers represent the minimum and maximum values.

**Figure supplement 5.**
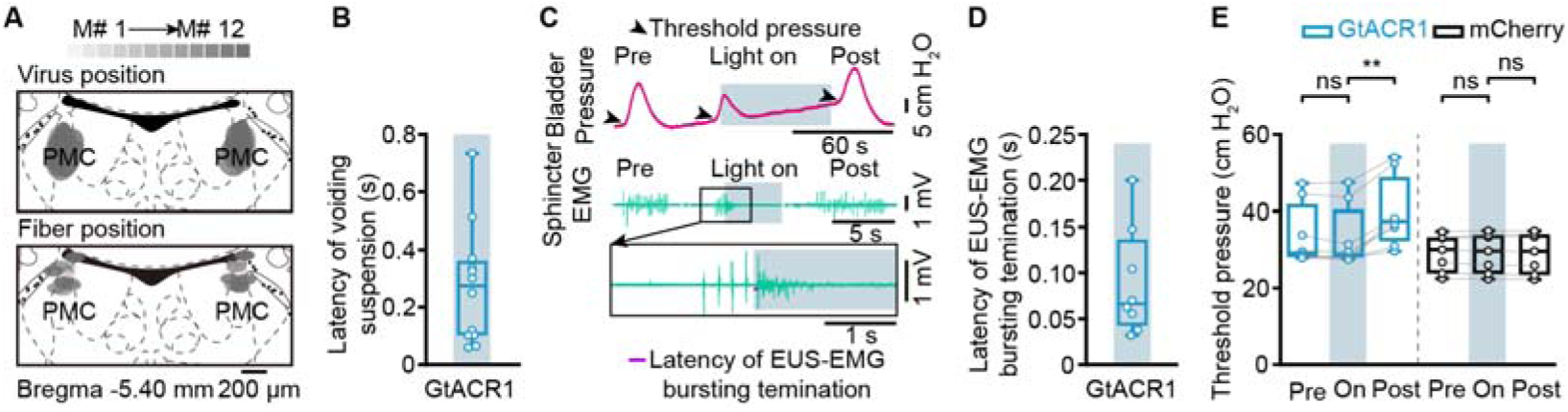
Acute photoinhibition (60 s) of PMC cells operates the bladder and sphincter to suspend voiding, related to Figure 2. (A) Overlay of viral expression areas (top) and fiber positions (bottom) from ESR1-Cre mice labeled with GtACR1 (n = 12 mice). (B) Latency of voiding suspension after light activation. (C) Representative raw traces of bladder pressure and EUS-EMG. (D) Latency of sphincter bursting termination after light activation (n = 8 mice). (E) Bladder threshold pressure during voiding before (‘Pre’), during (‘On’), and after (‘Post’) photoinhibition in PMC^ESR1-GtACR1^ (n = 8 mice) and PMC^ESR1-mCherry^ (n = 7 mice) groups (from left to right: *P* = 0.06, \*\**P* = 7.8e-3, *P* = 0.9, *P* = 0.4, respectively; n.s., not significant; Wilcoxon signed-rank test). For all data points in (B, D) and (E), whisker-box plots indicate the median with the 25%-75% percentile as the box, and whiskers represent the minimum and maximum values.

**Figure supplement 6.**
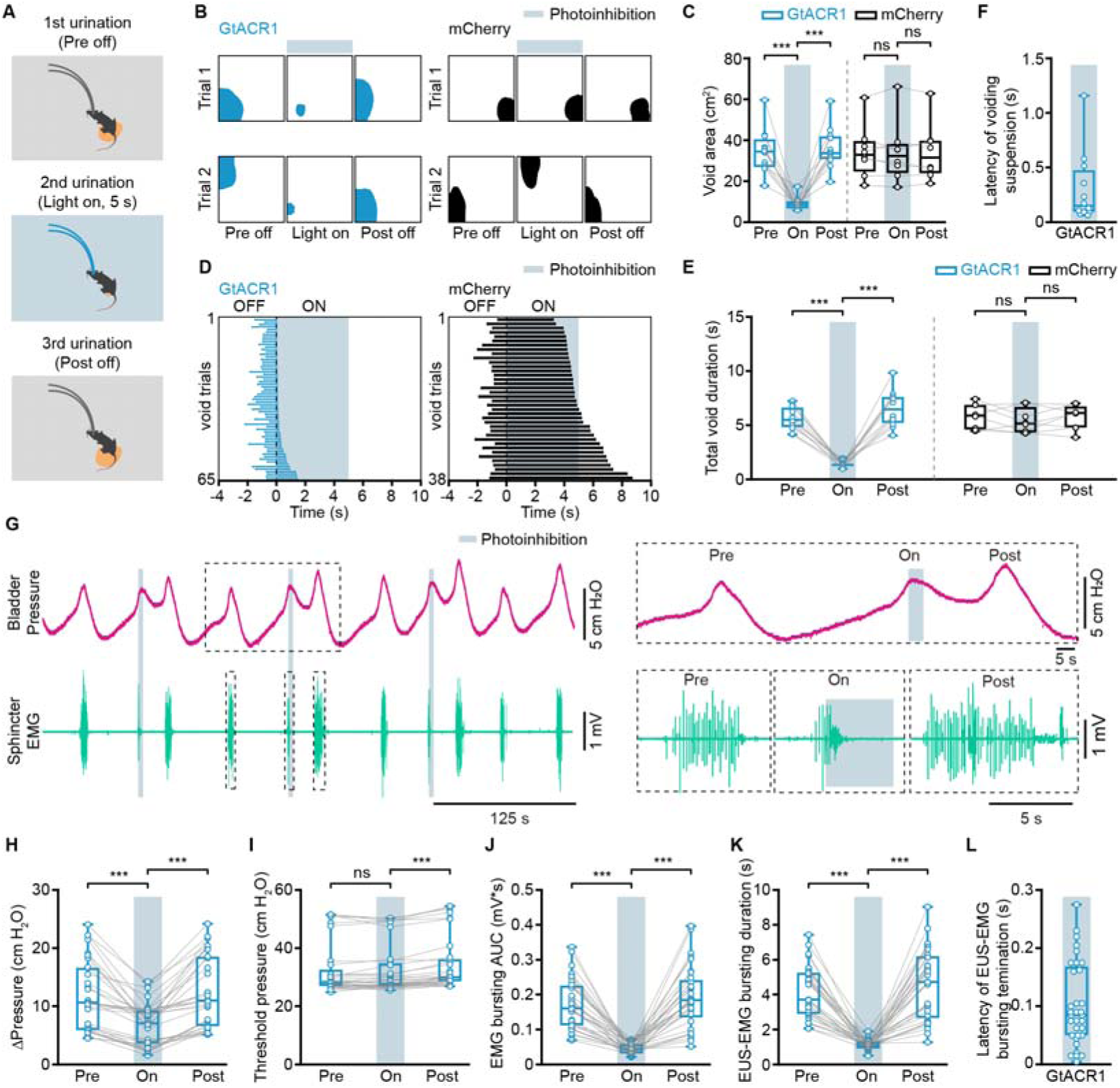
5-s photoinhibition of PMC cells suppresses bladder contraction and EUS bursting activity to suspend ongoing voiding. (A) Experimental design for 5 s photoinhibition of PMC^ESR1+^ cells in a freely moving mouse. (B, C) Representative images (B, blue and black shading) and quantification (C) of the void area before, during, and after 5 s photoinhibition in PMC^ESR1-GtACR1^ (n = 12 mice) and PMC^ESR1-mCherry^ (n = 8 mice) groups (from left to right: \*\*\**P* = 4.9e-4, \*\*\**P* = 4.9e-4, *P* = 0.7, *P* = 0.9, respectively; n.s., not significant; Wilcoxon signed-rank test). (D) Cumulative trials of voiding duration during 5 s photoinhibition in PMC^ESR1-GtACR1^ (blue bar, n = 65 trials from 12 mice) and PMC^ESR1-mCherry^ groups (black bar, n = 38 trials from 8 mice), ordered by increasing voiding epoch time with the laser on. (E) Voiding duration before, during, and after 5 s photoinhibition in PMC^ESR1-GtACR1^ (n = 12 mice) and PMC^ESR1-mCherry^ (n = 8 mice) groups (from left to right: \*\*\**P* = 4.9e-4, \*\*\**P* = 4.9e-4, *P* = 0.3, *P* = 0.4, respectively; n.s., not significant; Wilcoxon signed-rank test). (F) Latency of urination suspension after light activation. (G) Representative traces (left) and expanded portions (right, from the dashed box in the left panel) of bladder pressure (magenta) and EUS-EMG (teal) before, during, and after 5 s photoinhibition in PMC^ESR1-GtACR1^ individual. (H-L) Quantification of the effect of 5 s photoinhibition (n = 33 trials from 6 mice) on bladder pressure and EUS-EMG: Δpressure (H, \*\*\**P* = 5.4e-7), threshold pressure (I, *P* = 0.98, \*\*\**P* = 1.9e-6, respectively), EUS-EMG bursting AUC (J, \*\*\**P* = 5.4e-7), and EUS-EMG bursting duration (K, \*\*\**P* = 5.4e-7; n.s., not significant; Wilcoxon signed-rank test). (L) Latency of sphincter bursting termination after 5 s photoinhibition. For all data points in (C), (E), (F), and (H-L), whisker-box plots indicate the median with the 25%-75% percentile as the box, and whiskers represent the minimum and maximum values.

**Figure supplement 7.**
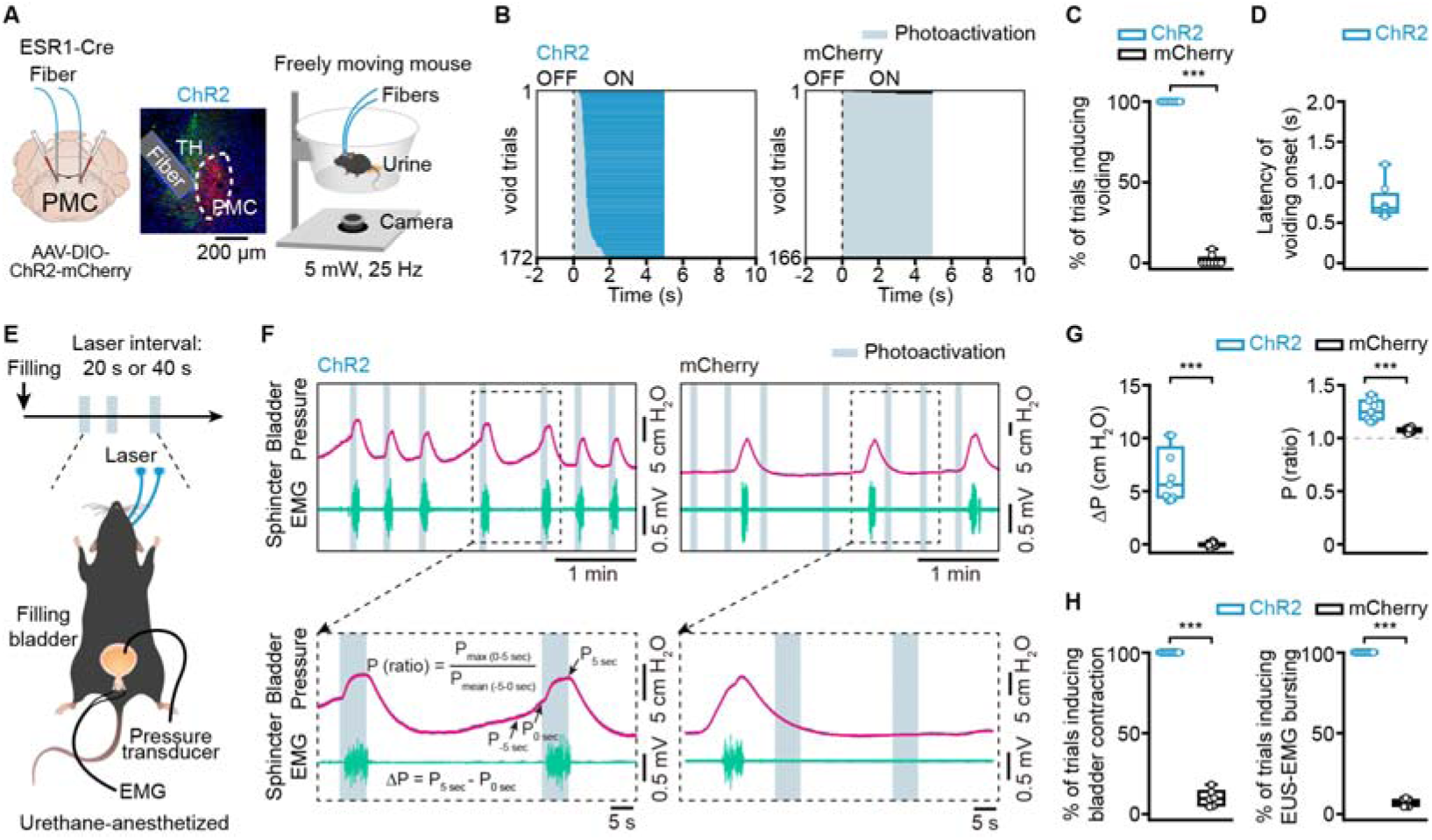
Activation of PMC cells induces both bladder contraction and EUS bursting activity to initiate voiding. (A) Schematics of labeling (left), representative histology (middle), and behavior test (right) for PMC^ESR1+^ photoactivation. Scale bar: 200 µm. (B) Cumulative trials of voiding duration in PMC^ESR1-ChR2^ (blue bar, n = 172 trials from 8 mice) and PMC^ESR1-mCherry^ (black bar, n = 166 trials from 8 mice) photoactivation. Voiding trials are ordered by the latency of the voiding epoch with the laser on. (C) % of photoactivation-associated voiding events in PMC^ESR1-ChR2^ and PMC^ESR1-mCherry^ groups (n = 8 mice per group, \*\*\**P* = 1.6e-4, Wilcoxon rank-sum test). (D) Latency of voiding onset after light on. (E) Timeline (top) and schematics (bottom) for PMC^ESR1+^ cells photoactivation during simultaneous cystometry and electromyography recording. (F) Representative traces (top) and expanded portions (bottom) of bladder pressure (magenta) and EUS-EMG (teal) around photoactivation timepoint in PMC^ESR1-ChR2^(left) or PMC^ESR1-mCherry^ (right) groups. (G, H) Quantification of bladder pressure change (ΔP, G, left), the ratio of bladder pressure (G, right), the percentage of photoactivation-associated bladder contraction (H, left), and the percentage of photoactivation-associated EUS-EMG bursting (H, right) upon photoactivation (n = 9 PMC^ESR1-ChR2^ mice, n = 6 PMC^ESR1-mCherry^ mice, \*\*\**P* = 4e-4 for (G, H), Wilcoxon rank-sum test). For all data points in (C, D, G), and (H), whisker-box plots indicate the median with the 25%-75% percentile as the box, and whiskers represent the minimum and maximum values.

**Figure supplement 8.**
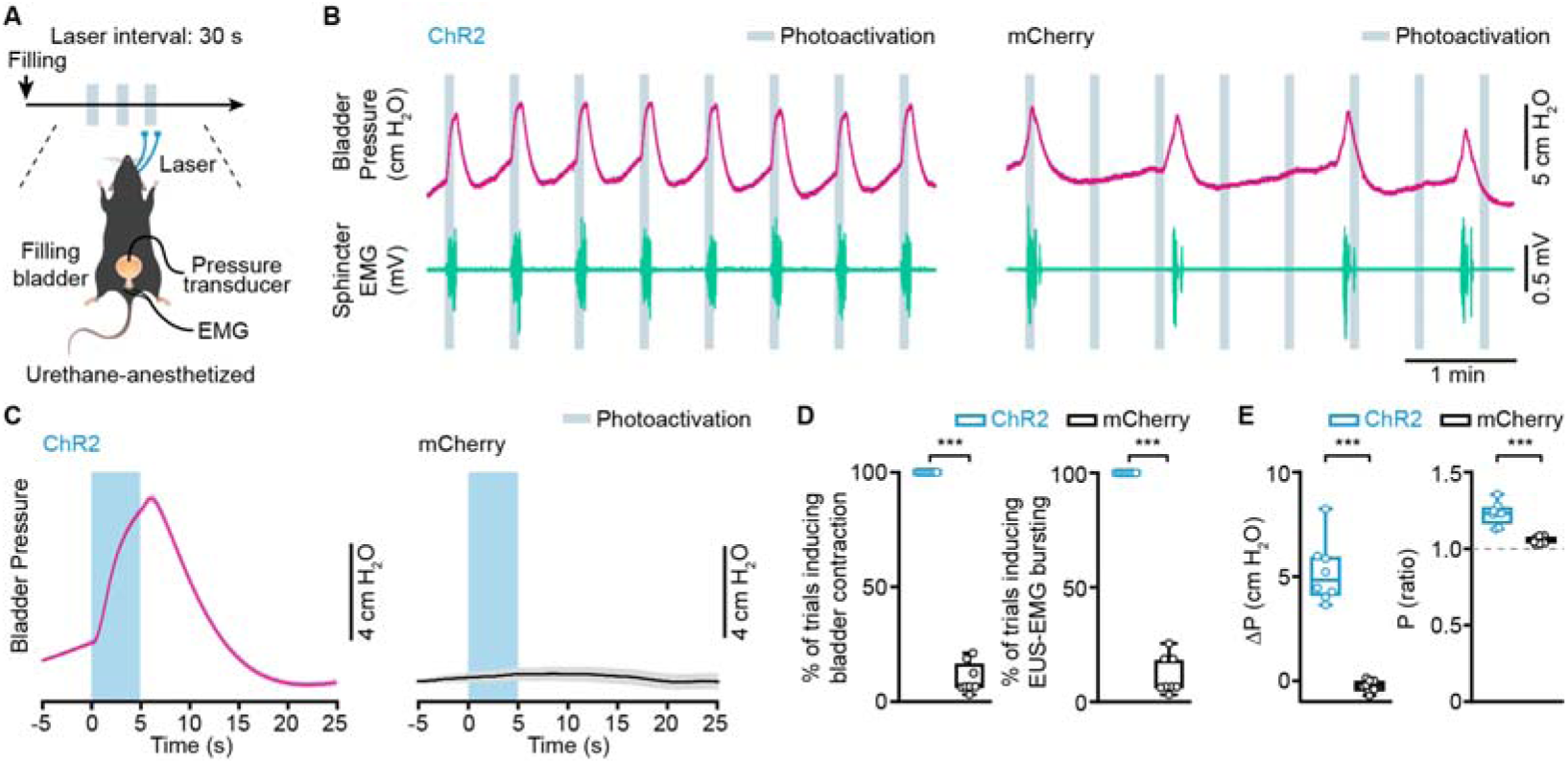
Regular interval photoactivation of PMC cells induces both bladder contraction and EUS bursting activity. (A) Timeline (top) and schematic (bottom) for regular interval photoactivation of PMC^ESR1+^ cells during simultaneous cystometry and urethral electromyography recording with a filled bladder. (B) Representative raw traces of bladder pressure (magenta) and EUS-EMG (teal) around the photoactivation timepoint in PMC^ESR1-ChR2^ (left) and PMC^ESR1-mCherry^ (right) individuals. (C) Average bladder pressure around photoactivation in PMC^ESR1-ChR2^ (left, n = 8 mice) and PMC^ESR1-mCherry^ (right, n = 8 mice) groups. The thick line and shading represent mean ± s.e.m., respectively. (D, E) Quantification of the photoactivation effect on the bladder detrusor and urethral sphincter in PMC^ESR1-ChR2^ (n = 8 mice) or PMC^ESR1-mCherry^ (n = 8 mice) groups: the percentage of bladder contraction (D, left), the percentage of EUS-EMG bursting (D, right), Δpressure (E, left), and bladder pressure ratio (E, right; \*\*\**P* = 1.6e-4 for D and E, Wilcoxon rank-sum test). For all data points in (D, E), whisker-box plots indicate the median with the 25%-75% percentile as the box, and whiskers represent the minimum and maximum values.

**Figure supplement 9.**
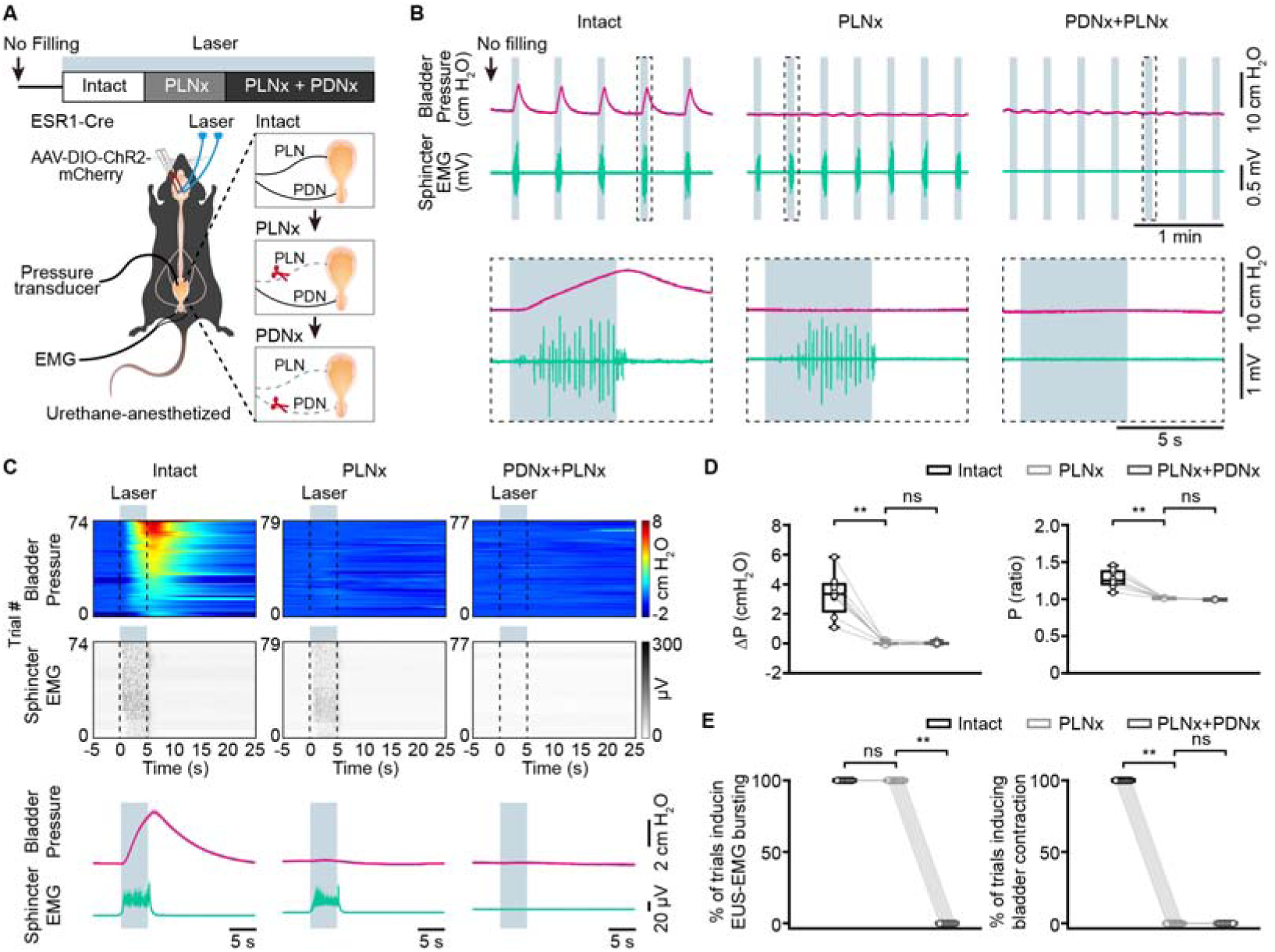
Transection of the pelvic nerves does not alter bladder pressure in an unfilled bladder during PMC^ESR1-ChR2^ photoactivation. (A) Timeline (top) and schematic (bottom) for PMC^ESR1-ChR2^ photoactivation during simultaneous cystometry and urethral electromyography recordings in a non-filled bladder, with PLNX performed first. PLNx: pelvic nerve transection; PDNx: pudendal nerve transection. (B) Representative traces (top) and expanded portions (bottom, from the dashed box in the top panel) showing bladder pressure (magenta) and EUS-EMG (teal) during PMC^ESR1-ChR2^ photoactivation in an unfilled bladder with PLNX performed first. (C) Heatmap (top) and average traces (bottom, thick line, and shading represent mean ± s.e.m., respectively) of sorted bladder pressure and EUS-EMG around photoactivation timepoint for all unfilled bladder trials with PLNX performed first (n = 8 mice per group). (D, E) Quantification of bladder pressure change (ΔP, D, left), bladder pressure ratio (D, right), the percentage of photoactivation-associated EUS-EMG bursting (E, left), and the percentage of photoactivation-associated bladder contraction (E, right) upon photoactivation for the PLNx-first experiment from C (n = 8 mice per group, from left to right, D: \*\**P* = 7.8e-3, *P* =0.8, \*\**P* = 7.8e-3, *P* = 0.1, respectively; E: *P* = 1, \*\**P* = 7.8e-3, \*\**P* = 7.8e-3, *P* = 1, respectively; n.s., not significant; Wilcoxon signed-rank test). For all data points in (D, E) whisker-box plots indicate the median with the 25%-75% percentile as the box, and whiskers represent the minimum and maximum values.

**Figure supplement 10.**
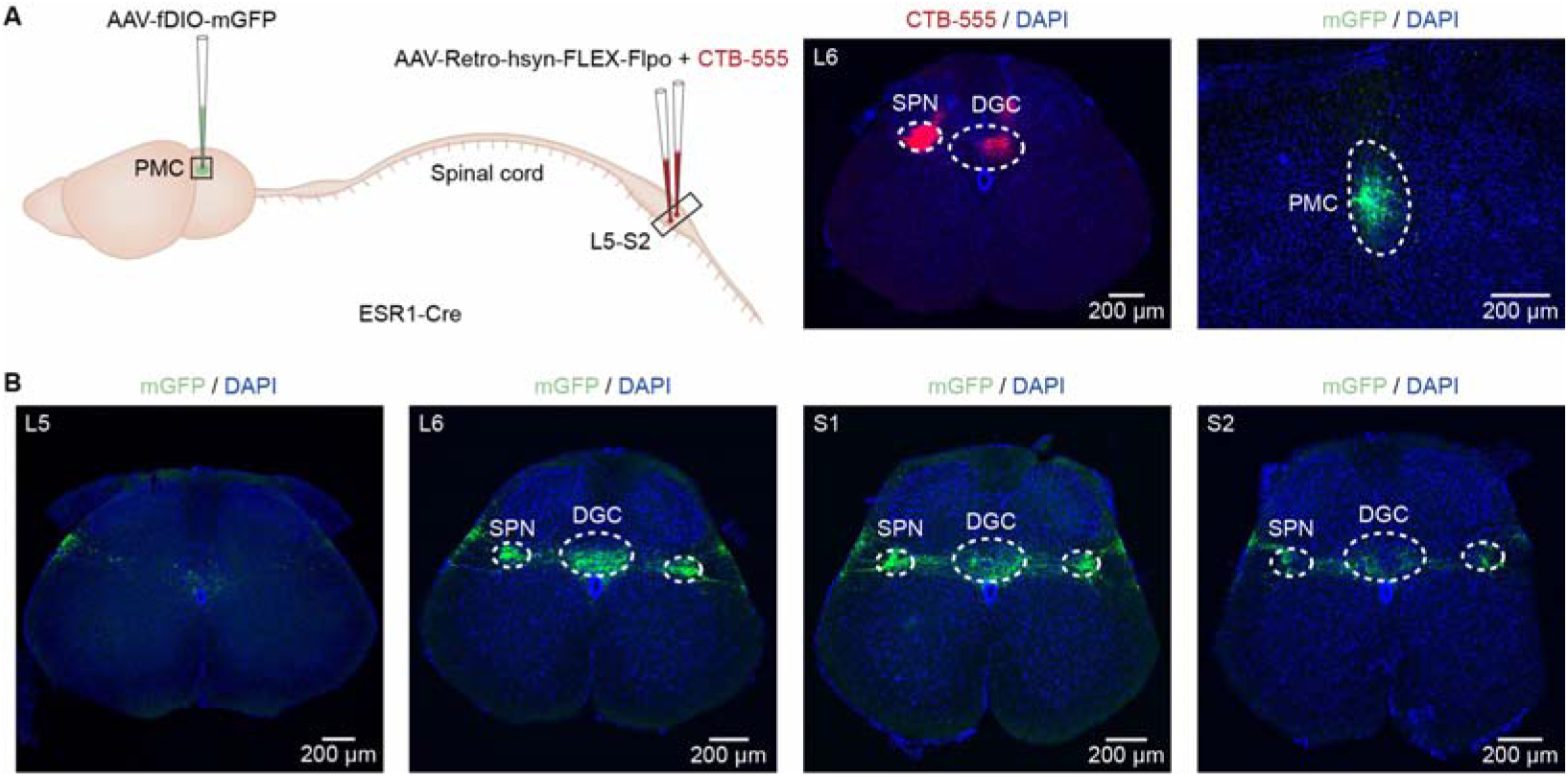
PMC cells projection to the SPN and DGC in the spinal cord. (A) Left: Schematic of labeling. Middle and right: Representative histological images showing CTB-555 expression in the lumbosacral spinal cord (middle) and mGFP expression in the PMC (right). Scale bars: 200 µm. (B) Axonal projections of PMC^ESR1-mGFP^ cells in the lumbosacral spinal cord from L5 to S2 levels (n = 3 mice). Scale bars: 200 µm. Abbreviations: SPN, sacral parasympathetic nucleus; DGC, dorsal gray commissure.

**Figure supplement 11.**
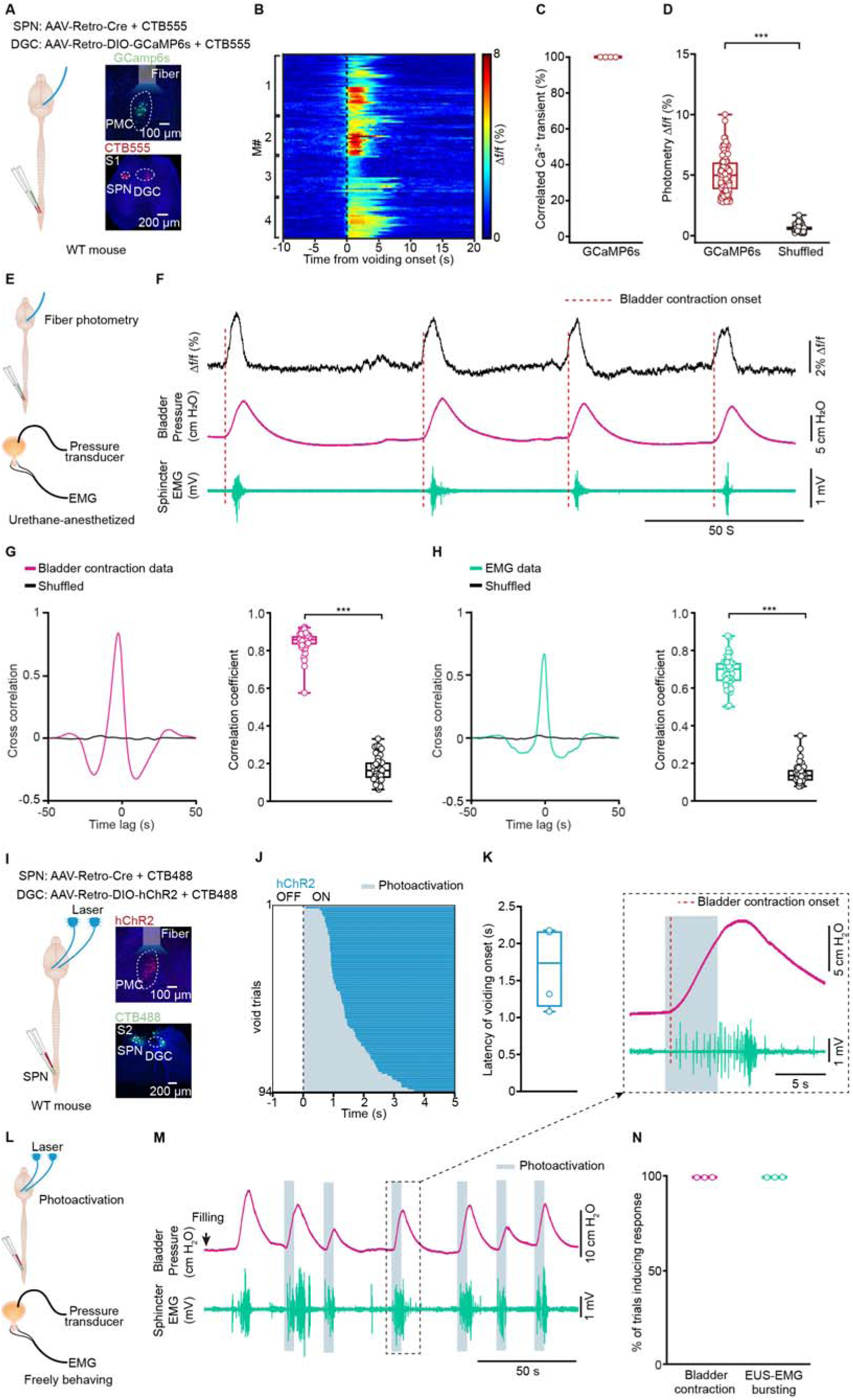
Dual-projecting PMC neurons are active throughout voiding, correlating with bladder pressure and EUS bursting, and their optogenetic activation initiates urination. (A) Left: Schematic of the labeling strategy. Right: Representative histological images showing GCaMP6s expression in the PMC and CTB-555 labeling in the lumbosacral spinal cord of a wild-type mouse. Scale bars: 100 µm (top), 200 µm (bottom). (B) Cumulative sessions of Ca^2+^ signals aligned to voiding onset (dotted line). (C) Quantification of voiding-related Ca^2+^ events from 4 mice. (D) Boxplots showing the amplitude of voiding-related Ca^2+^ signals in various groups (n = 102 trials from 4 mice per group, 5.0%\3.8%-6.0% for GCaMP6s, 0.6%\0.4%-0.8% for shuffled, \*\*\**P* = 1.8e-18, Wilcoxon signed-rank test). (E) Schematic of fiber photometry recording for dual-projecting PMC cells during simultaneous cystometry and EUS-EMG in a urethane-anesthetized mouse. (F) Representative traces showing Ca^2+^ transients (black), bladder pressure (magenta), and EUS-EMG (teal) during fiber photometry recordings. Dashed lines indicate the onset of bladder contractions. (G) Cross-correlation (left) and correlation coefficients (right) between Ca^2+^ signals and bladder contraction events, compared to shuffled data (n = 40 trials from 4 mice, \*\*\**P* = 3.6e-8, Wilcoxon signed-rank test). (H) Cross-correlation (left) and correlation coefficients (right) between Ca^2+^ signals and EUS-EMG bursting events, compared to shuffled data (n = 40 trials from 4 mice, \*\*\**P* = 3.6e-8, Wilcoxon signed-rank test). (I) Left: Schematic of the labeling strategy. Right: Representative histological images showing hChR2 expression in the PMC and CTB-488 labeling in the lumbosacral spinal cord of a wild-type mouse. Scale bars: 100 µm (top), 200 µm (bottom). (J) Cumulative trials of voiding duration in dual-projecting PMC cells photoactivation (blue bar, n = 94 trials from 4 mice). Voiding trials are ordered by the latency of the voiding epoch with the laser on. (K) Latency of voiding onset after light on. (L) Schematic for dual-projecting PMC cells photoactivation during simultaneous cystometry and EUS-EMG in a freely behaving mouse. (M) Representative traces and expanded portion (black dashed box) of bladder pressure (magenta) and EUS-EMG (teal) around photoactivation timepoint. Dashed line indicates bladder contraction onset. (N) The percentage of photoactivation-associated bladder contraction and EUS-EMG bursting activity (n = 3 mice per group). For all data points in (C, D, G, H, K), and (N), whisker-box plots indicate the median with the 25%-75% percentile as the box, and whiskers represent the minimum and maximum values.

**Figure supplement 12.**
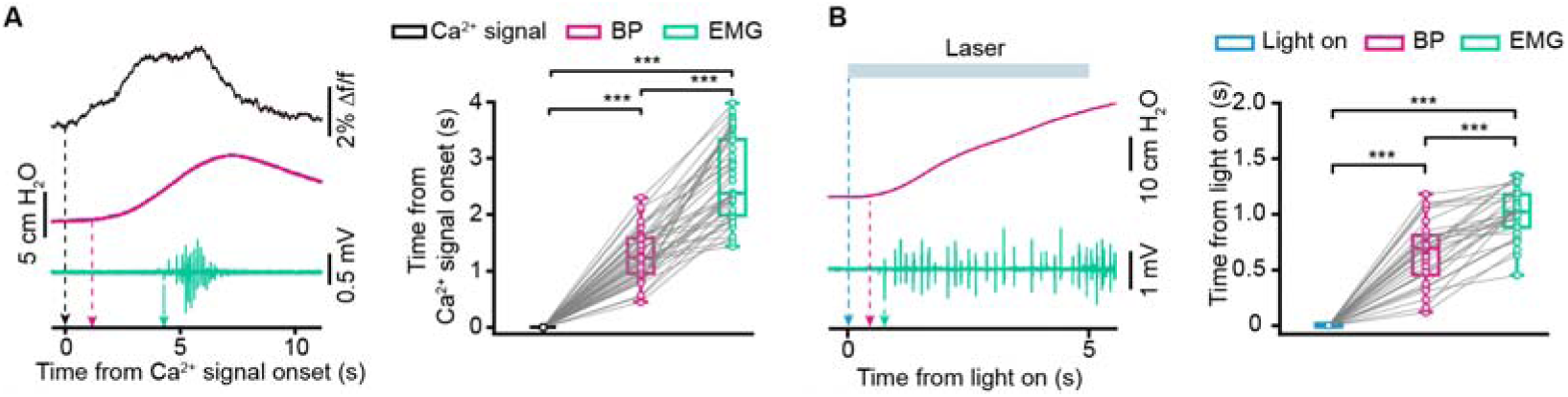
Dual-projecting PMC neurons coordinate bladder contraction and EUS bursting activity. (A) Left: Example (left, with arrows indicating the onset timepoints) and quantification (right) of the temporal relationships among the onset times of Ca^2+^ signals, bladder pressure (BP) upstroke, and EUS-EMG bursting in photometry recordings (n = 40 trials from 4 mice; Ca^2+^signals onset, 0\0-0 s; BP upstroke onset, 1.2\0.9-1.6 s; EUS-EMG bursting onset, 2.3\2.0-3.3 s; \*\*\**P* = 3.6e-8 for all, Wilcoxon signed-rank test). (B) Example (left, with arrows indicating the onset timepoints) and quantification (right) of the temporal relationships among the onset times of light stimulation, bladder pressure upstroke, and EUS-EMG bursting for the photoactivation group (n = 30 trials from 3 mice; Light on, 0\0-0 s; BP upstroke onset, 0.7\0.5-0.8 s; EUS-EMG bursting onset, 1.0\0.9-1.2 s; \*\*\**P* = 1.7e-6 for all, Wilcoxon signed-rank test). For all data points in (A, B), whisker-box plots indicate the median with the 25%-75% percentile as the box, and whiskers represent the minimum and maximum values.

